# From deep TLS validation to ensembles of atomic models built from elemental motions

**DOI:** 10.1101/012930

**Authors:** Alexre Urzhumtsev, Pavel V. Afonine, Andrew H. Van Benschoten, James S. Fraser, Paul D. Adams

## Abstract

The widely used *Translation Libration Screw* (TLS) approximation describes concerted motions of atomic groups in X-ray refinement. TLS refinement often provides a better interpretation of diffraction data and the resulting rigid body motions may subsequently be assigned biochemical significance. In TLS refinement, three matrices (*T, L and S*) describe harmonic vibration, libration and their correlation. Because these matrices describe specific motions, they impose a number of conditions on their elements. Ignoring these conditions while refining the matrix elements may result in matrices that cannot be interpreted in terms of physically realistic motions. We describe a mathematical framework and the computational tools to analyze refined *TLS* matrices through their decomposition into descriptors of underlying motions. This allows for straightforward validation and identification of implausible *TLS* parameters. An algorithm for the generation of structural ensembles that are consistent with given *TLS* parameters, implemented as a part of the *Phenix* project, is also described.

**Synopsis:** Procedures for the decomposition of *TLS* matrices into elementary vibrations and librations indicates possible errors in the definition of these matrices and corrects them when possible. The program outputs the corresponding vibration-libration parameters and generates structural ensembles.

## 1. Introduction

Crystallographic models fit to X-ray, neutron, or electron diffraction data describe each atom by its central position **r_0_** and additional parameters. Structural disorder, particularly thermal motion, is primarily fit by the so-called Debye-Waller factor (also known as the B-factor or displacement parameter) that reflects the probability of an atom moving from its central position. This factor is accounted for as a multiplier for each structure factor corresponding to integer indices (*h*, *k*, *l*). In a harmonic approximation, this factor can be presented as following (see for example, Grosse-Kunstleve & Adams, 2002, and references therein)

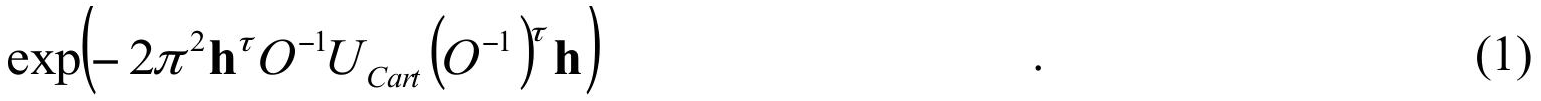

Here *O* is the orthogonalization matrix for the given crystal, **h** is the column vector with the indices (*h*, *k*, *l*), and *τ* signifies matrix or vector transposition. (In Grosse-Kunstleve & Adams (2002) the orthogonalization matrix is noted as *A*; here this letter is used for the matrix in development of *U*_*Cart*_, following Tickle & Moss, 1999 and Urzhumtsev et al., 2013). The symmetric matrix *U*_*Cart*_ is defined by the average shifts (and their correlations) along the coordinate axes. The matrix *U*_*Cart*_ varies between atoms and for isotropic atoms is diagonal with equal elements.

However, purely isotropic motion is rarely observed in macromolecular crystals where the displacement of groups of atoms can be correlated. For example, while each atom of a side chain may oscillate independently around its central position, the side chain itself may oscillate around one (or several) bonds. This concerted motion is of special interest for several reasons. First, it may characterize the mobility of macromolecular domains and thus give insight into molecular mechanism. Second, describing only the common motion may drastically simplify the model and reduce the number of parameters needed to model the data. This may be especially attractive when working at relatively low resolutions.

A description of concerted molecular motions has been introduced by Cruickshank (1956) and Schomaker & Trueblood (1968) and later developed in a number of works (for example Johnson, 1970; Scheringer, 1973; Howlin et al., 1989, 1993; Kuriyan & Weis, 1991; Schomaker & Trueblood, 1998; Tickle & Moss, 1999; Winn et al., 2001, 2003; Painter & Merritt, 2005, 2006a, 2006b; for a recent comprehensive review see Urzhumtsev et al. 2013). Since rigid body displacement is a composition of translation and rotation (see for example Goldstein, 1950), Schomaker & Trueblood presented the matrices *U*_*cart,n*_ for all atoms *n* = 1, 2, … *N* within the model as a sum

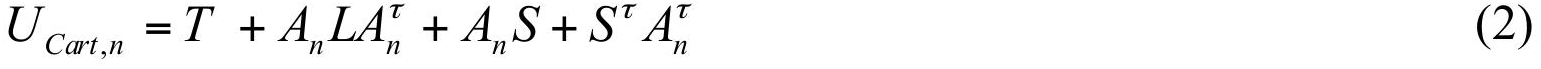

Anti-symmetric matrices *A*_*n*_ are functions of the Cartesian coordinates (*x*_*n*_, *y*_*n*_, *z*_*n*_) of atom *n*

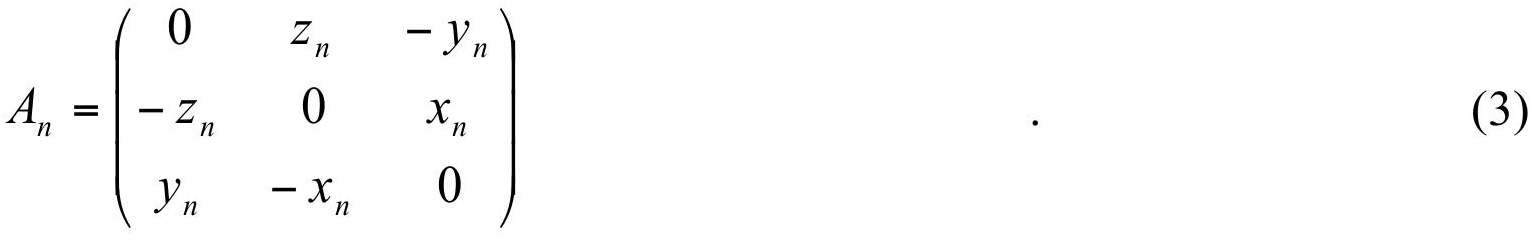

Matrix *S* and symmetric matrices *T* and *L* are common to all atoms of the moving group and are defined by the following elementary motions. *L* describes libration (oscillating rotations) around three principal rotation axes mutually orthogonal to each other. *T* describes an apparent translation of the atomic group (a term ‘vibration’ may be more appropriate for random translations around a central position). *S* describes the correlation between librations and vibrations (for a detailed definition Urzhumtsev et al. 2013). We use the term ‘apparent translation’ because matrix *T* may have an additional contribution from librations as discussed in Section 2.

Compared to calculating *TLS* matrices from corresponding libration and vibration parameters, the inverse procedure of decomposing TLS matrices into these parameters is more difficult. As discussed previously (for example, Johnson, 1970; Scheringer, 1973; Tickle & Moss, 1999) the problem itself is poorly posed since the same set of diffraction data (and as a consequence the same set of the *TLS* matrices) may correspond to different common motions of the contributing atoms or atomic groups. Nevertheless, TLS refinement allows one to search for composite motions (librations and vibrations) that correspond to a particular set of matrices. Thus, one may refine either the parameters of the composite motions or the elements of the *TLS* matrices as independent parameters, as is currently implemented in modern refinement programs,.

To describe underlying elementary motions, the matrices must obey certain conditions; otherwise, the physical motions they represent are nonsensical. First, there is the trivial condition that the matrices *T* and *L* must be symmetric and positive semidefinite. Second, the matrices must be compatible with each other. For example, since matrix *S* describes the correlation between libration and vibration, its elements should be consistent with the elements of *T* and *L*. Ignoring this condition may result in a non-positive definite matrix *T* after the apparent contribution of libration is subtracted. Interestingly, refinement of *TLS* matrix elements without taking into account these conditions can still result in physically reasonable individual atomic displacement parameters and an improved model to data fit. Zucker et al. (2010) analyzed all PDB entries (Bernstein *et al*., 1977; Berman *et al*., 2000) that employ *TLS* and suggested tools for validating *TLS* parameters to ensure that individual atomic displacement parameters were physically realistic. While programs for the extraction of libration and vibration parameters from *TLS* matrices (Howlin et al., 1993; Painter & Merritt, 2005). A comprehensive mathematical protocol for accomplishing this task could be helpful for determining if decomposition into physically interpretable motions is even possible. This manuscript describes the corresponding algorithm (Fig. 1) and provides practical calculation protocols in a ready-to-program style. In particular, we discuss a list of conditions defining the plausibility of such decomposition. Below, we discuss the decomposition of a single set of *TLS* matrices; however, the procedure is similar when molecular models are composed of several such groups. For a PDB model that contain several sets of *TLS* matrices, each set can be processed independently.

**Figure 1.**
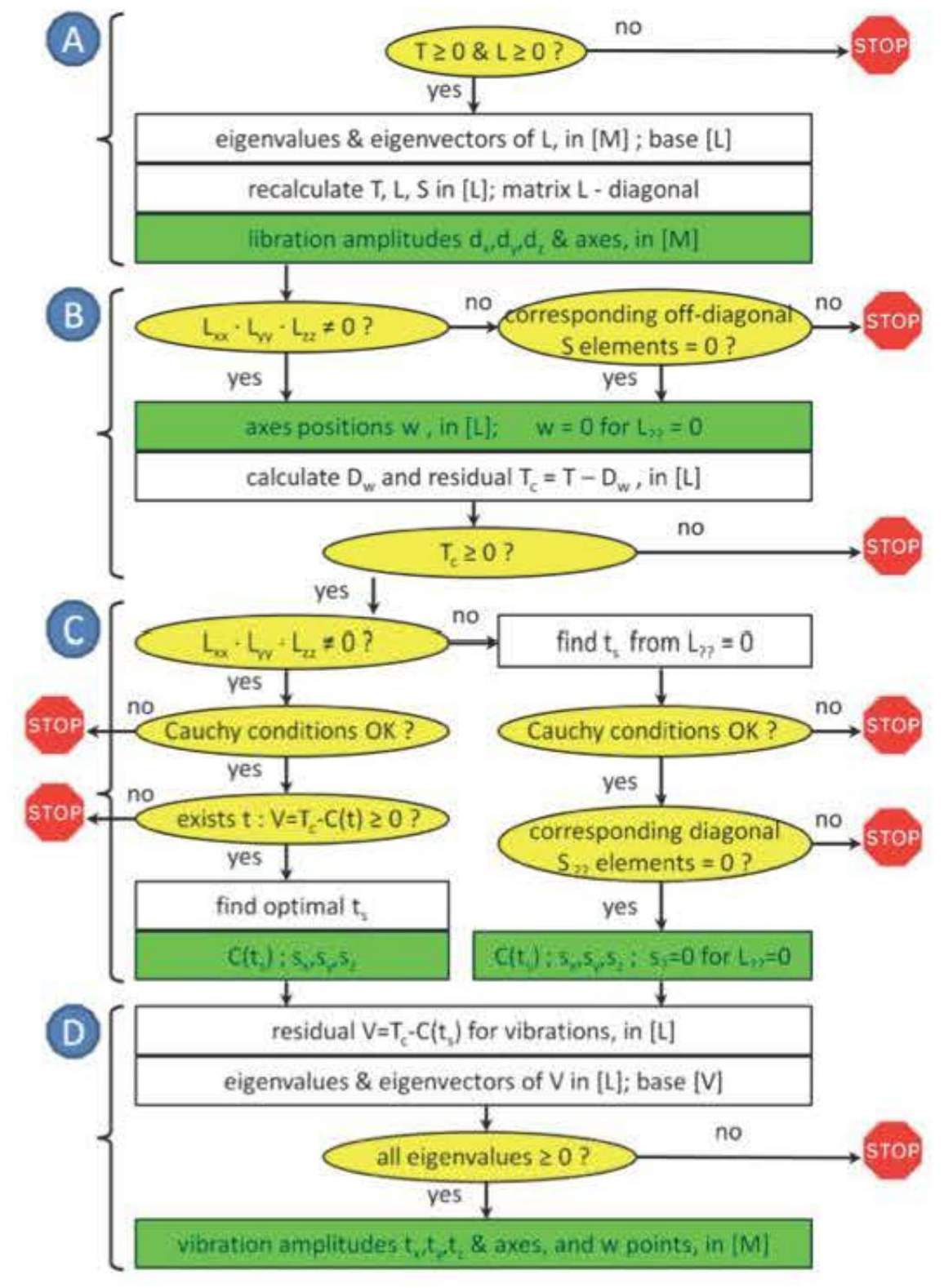
General flowchart of the *TLS* decomposition into libration and vibration composite motions. Yellow ellipses are for conditions to the verified. Green rectangles are for the output parameters of the composite motions. Letters A-E indicate different steps of the procedure as described in the text.

Thus, analyzing the physical reasonableness of the individual atomic displacement parameters and the rigid-body motions that give rise to them require separate validation tools. Indeed, from roughly 200,000 sets of *TLS* matrices in the PDB (25,284 entries from the total 194,633 PDB depositions) about one-third contain *T* or *L* matrices that are non positive-semidefinite and another one-third (Table 1) cannot describe vibration-libration correlated motions. Although these numbers can be slightly reduced by removing marginal cases, these results suggest that decomposition of *TLS* matrices into elementary motions can be an important validation step. This analysis can be used to build ensembles of models representing plausible conformations, which will allow for a better description of diffraction data and for the testing of model agreement with diffuse X-ray scattering (Van Benschoten *et al*., accompanying paper).

**Table 1.** Number of PDB entries with at least one unphysical TLS matrix condition. Note:when one of the conditions was found to be broken, the other conditions were not checked (Figure 1). The conditions are (from left to right): (*T*≥0 & *L*≥0) matrices *T* and *L* are positive semidefinite; (s=0 & w=0) for the *L* matrix with one degenerate libration the corresponding elements of the *S* matrix are equal to 0; (*S*≤*TL*) elements of the *S* matrix are limited by the corresponding elements of the *T* and *L* matrices following the Cauchy conditions (23); (*TC*≥0) matrix *T* is positive semidefinite after the contribution due to the displacement of libration axes is removed; (*V*≥0) residual *V* matrix is positive semidefinite. Right columns summarize the total number of the *TLS* sets with one of the condition broken and the number of the PDB entries concerned. The two lines shows the statistics for the original matrices in the PDB with the default condition *trS* = 0 (upper line) and after the optimal choice of the diagonal *S* elements whenever possible as described in Sections 3-4 (bottom line). The total number of the PDB entries is 104,633.

| | total PDB<br>number | total $TLS$<br>number | conditions broken | | | | | total $TLS$<br>broken | total PDB<br>broken |
| --- | --- | --- | --- | --- | --- | --- | --- | --- | --- |
| | | | $T \geq 0$ & $L \geq 0$ | $s=0$ & $w=0$ | $S \leq TL$ | $TC \geq 0$ | $V \geq 0$ | | |
| $t_S = 0$ | 25284 | 197920 | 66438 | 13824 | 1271 | 43934 | 7941 | 133408 | 22263 |
| $best\ t_S$ | 25284 | 197920 | 66438 | 13824 | 0 | 43090 | 5699 | 129051 | 21540 |

## 2. TLS matrices and different bases

### 2.1. Matrix presentations and decomposition difficulties

Even if a rigid body is involved in several simultaneous motions (supposing that the amplitudes of these motions are relatively small and the motions are harmonic), each motion can be described by a libration around three mutually orthogonal axes and by a vibration following three other mutually orthogonal axes (Urzhumtsev et al. 2013, and references therein). For example, if the principal vibration axes coincide with the coordinate axes, and 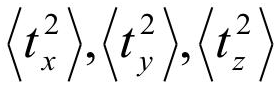 are the corresponding squared root-mean-square deviations (*rmsd*) along these axes, the matrix *T* in equation (2) is equal to

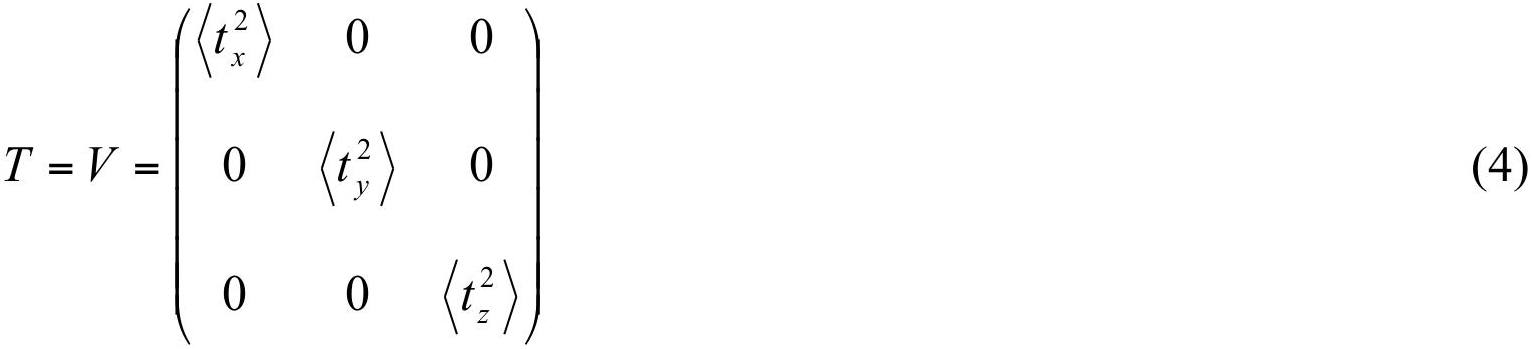

Similarly, if the libration axes coincide with the coordinate axes, the matrix *L* in (2) becomes:

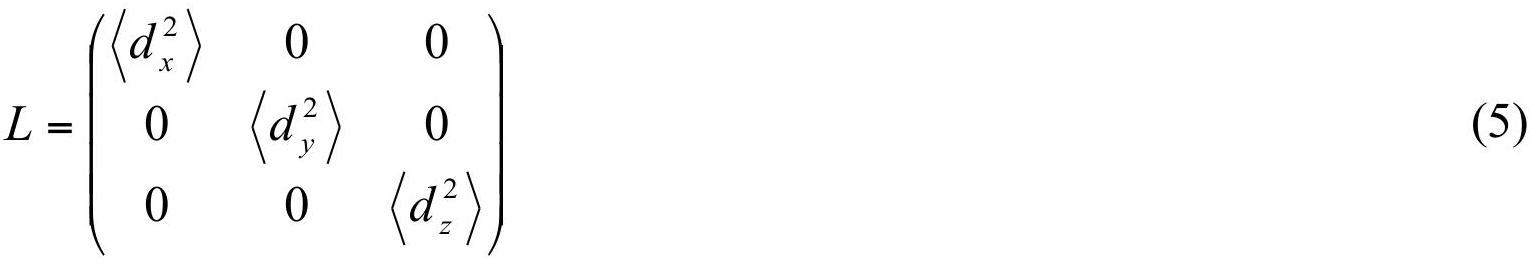

Here 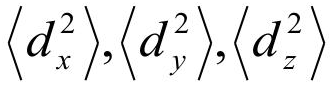 are the squared *rmsd*s of the vibration angle expressed in radians; for small deviations they are numerically equal to the squared *rmsd*s of points at a unit distance from the corresponding axes. One difficulty is that in practice the principal vibration and libration axes are not parallel to the coordinate axes, which requires transforming the matrices *T* and *L* into a more general form. Another difficulty is that vibration and libration motions may not be independent, as seen in helical rotations. If the correlations between translation and libration are characterized by parameters *s*_*x*_, *s*_*y*_, *s*_*z*_ (see Urzhumtsev et al. 2013, for a formal definition), the librations generate an additional contribution to translation so that matrix *T* becomes different from *V* as defined by equation (4) adding diagonal terms:

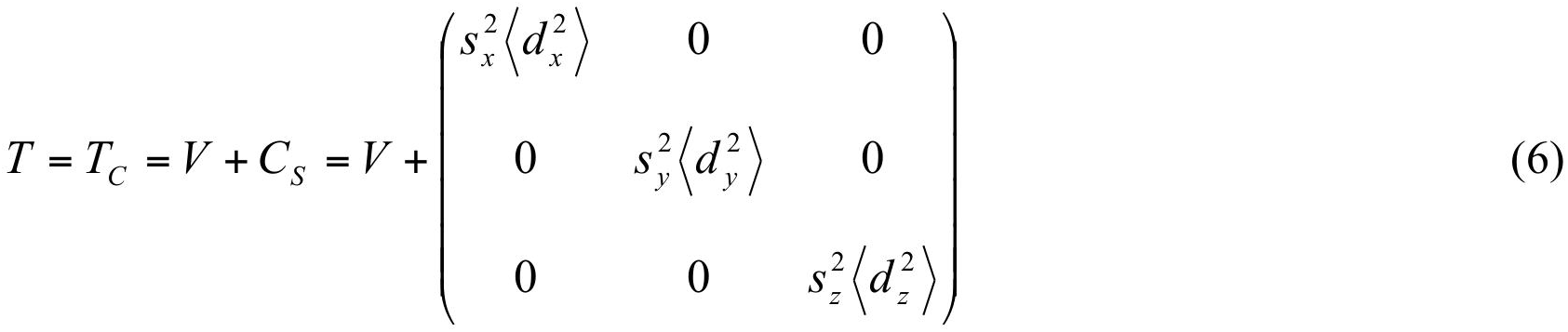

and also resulting in the non-zero *S* matrix

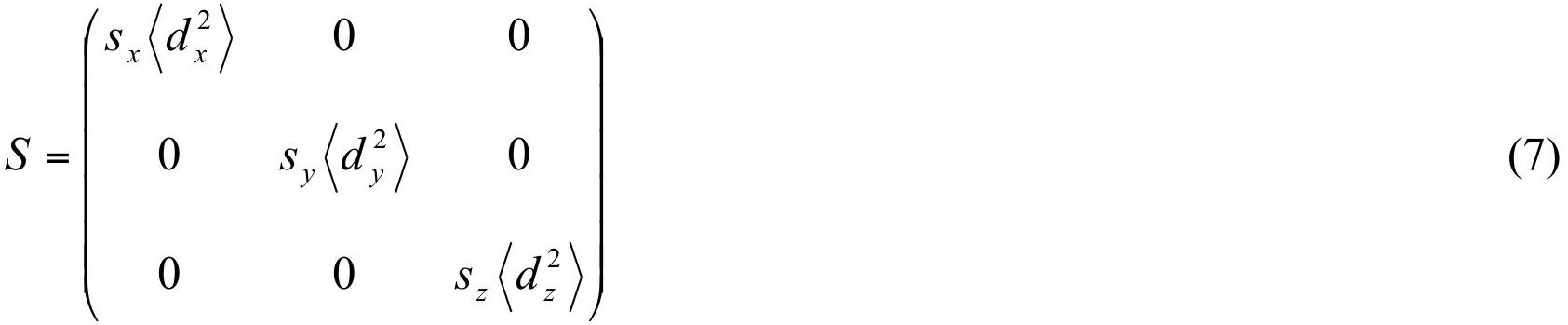

Yet another difficulty is that the principal libration axes do not need to pass through the origin, or even through a common point. This generates an additional apparent translation component and adds another term

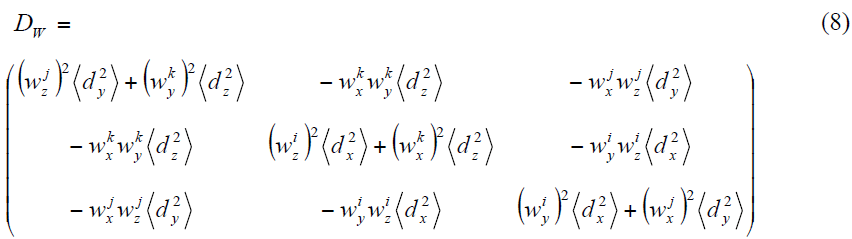

to the *T* matrix (Urzhumtsev et al. 2013). Similarly, matrix *S* must account for

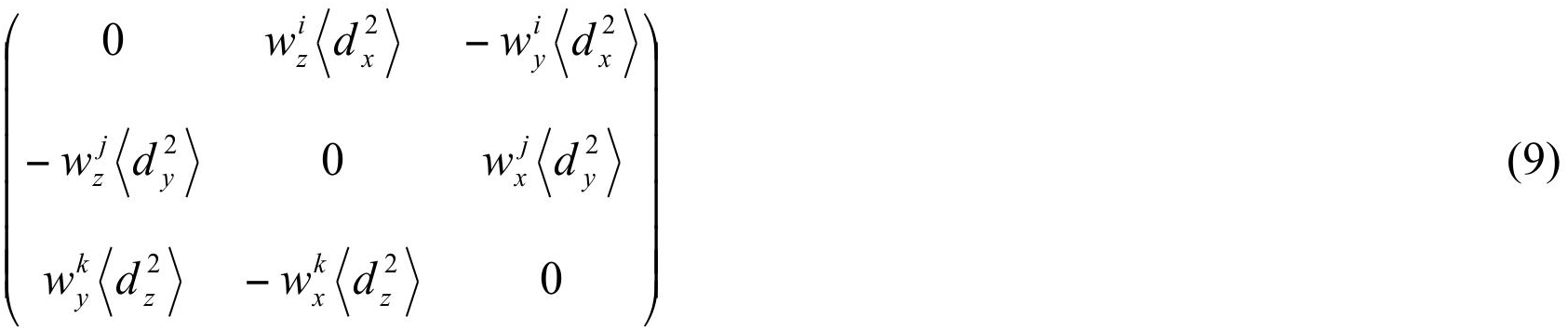

in addition to (7). Finally, the same actual motion can be presented by different combinations of the elementary motions since one may shift a rotation axis and compensate for this by an additional translation. The same total motion may also be split differently between correlated and independent contributions. These nuances complicate the procedure of extracting the descriptors of elementary motions from the *TLS* matrices.

### 2.2. Molecular basis and center of reaction

Formally, any origin (and, more generally, any basis) can be chosen to describe the composite motions. Clearly, description of the vibration does not depend on the choice of the origin, while the position of the libration axes changes as function of origin choice. Most often, the origin is taken in the center of mass of the atomic group or in between the axes so that the atomic displacements due to the choice of these axes are similar. This point is called the center of diffusion (Brenner, 1967) or the center of reaction (Tickle & Moss, 1999). Shifting from one origin to another does not change *L* but both *T*, minimizing its trace, and *S*, making it symmetric for the center of reaction (Brennen, 1967; Tickle & Moss, 1999; Urzhumtsev et al. 2013). Changing the origin does not modify the algorithm of the search for the composite motions. Thus, the *TLS* matrices depend on the choice of the coordinate basis and in particular its origin. In what follows, we suppose that the original *TLS* matrices are defined in a basis chosen with a relation to the molecule, which we will call the M basis (for ‘Molecular’). As we will be switching between different bases, we indicate the corresponding basis by index in brackets such as *T*_[*M*]_, *L*_[*M*]_ and *S*_[*M*]_.

## 3. Basis of libration axes

### 3.1. Diagonalization of the L matrix (L basis; step A)

We start the procedure from the matrix *L*_[*M*]_ that depends only on the libration parameters.

Let

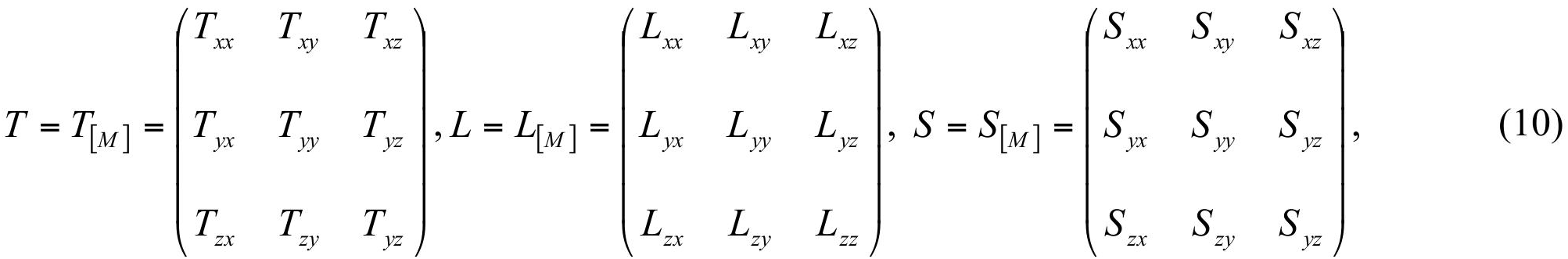

with *T*_*xy*_ = *T*_*yx*_, *T*_*xz*_ = *T*_*zx*_, *T*_*yz*_ = *T*_*zy*_ and *L*_*xy*_ = *L*_*yx*_, *L*_*xz*_ = *L*_*zx*_, *L*_*yz*_ = *L*_*zy*_, be the *TLS* matrices defined in the original M basis. Importantly, for further analysis of the *T* matrices, we remind the reader that if a matrix is symmetric, it remains symmetric for any rotation of the coordinate system.

The principal libration axes correspond to three mutually orthogonal eigenvectors of the symmetric matrix *L*_[*M*]_. First we search for the corresponding eigenvalues 0 ≤ *λ*_1_ ≤ *λ*_2_ ≤ *λ*_3_, which must be non-negative (see equation (5); eigenvalues do not change with the coordinate system). Let ***l*
_1_, *l*_2_, *l*_3_** be the corresponding normalized eigenvectors from which we construct a new basis*L*as

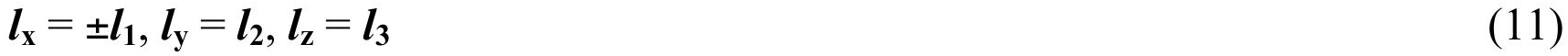

The appropriate sign for ***l***_**x**_ is chosen so that the vectors in (11) form a right-hand triad; for example one can take ***l***_**x**_ = ***l***_**y**_ × ***l***_**z**_ such that the condition is guaranteed.

The transition from the M basis into the L basis is a rotation with the corresponding matrix composed from the coordinates of the new basis vectors

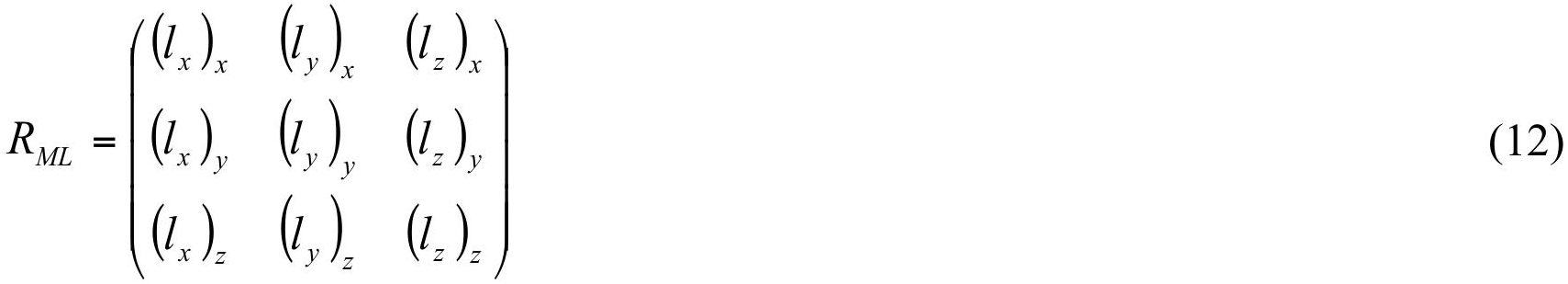

The *TLS* matrices in the L basis are

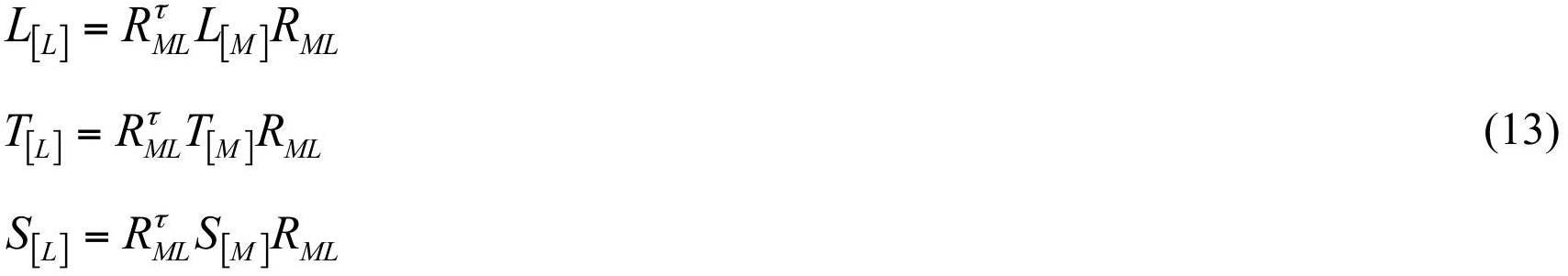

where matrix *L*_[*L*]_ is diagonal with the elements *L*_[*L*]*xx*_ = *λ*_1_*, L*_[*L*]*yy*_ = *λ*_2_*, L*_[*L*]*zz*_ = *λ*_3_ at its diagonal. The atomic coordinates in the L basis are

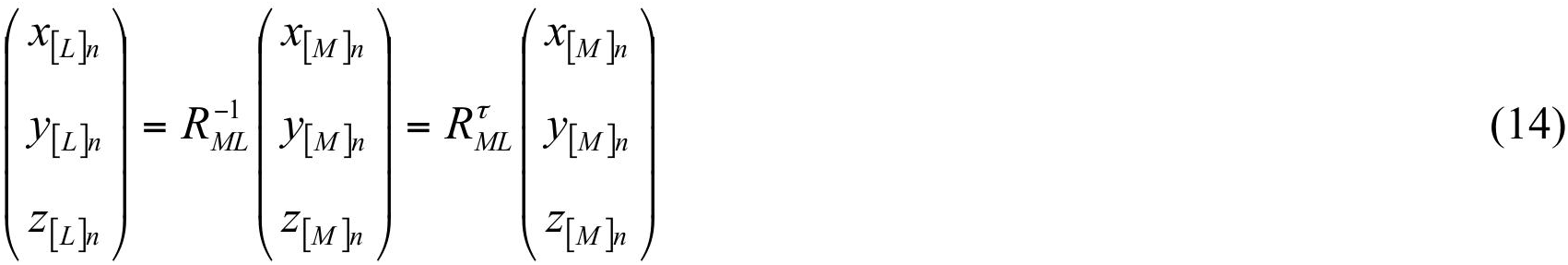

### 3.2. Position of the libration axes in the L basis (step B)

Given matrix *L*_[*L*]_ in the basis of its eigenvectors, we obtain the estimates 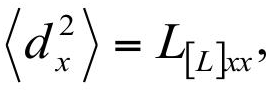
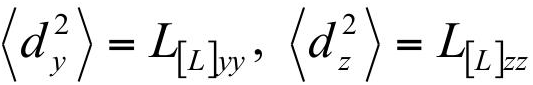 of the squared libration amplitudes around the three principal libration axes.

In the general case these libration axes do not pass through the origin but through some points 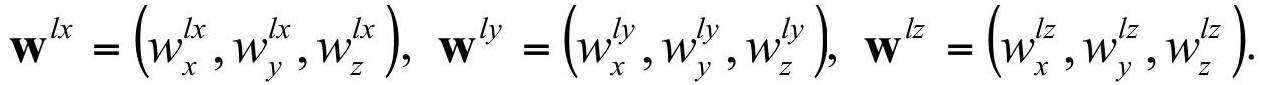 In fact, the *x*-component of **w**^*lx*^, *y*-component of **w**^*ly*^ and *z*-component of **w**^*lz*^ can be any values. In other words, the coordinate axes chosen by construction as eigenvectors of *L*_[*L*]_ are parallel to the libration axes but do not necessarily coincide with them.

Using equations (5) and (9) we calculate the positions of the rotation axes (Urzhumtsev et al. 2013) as

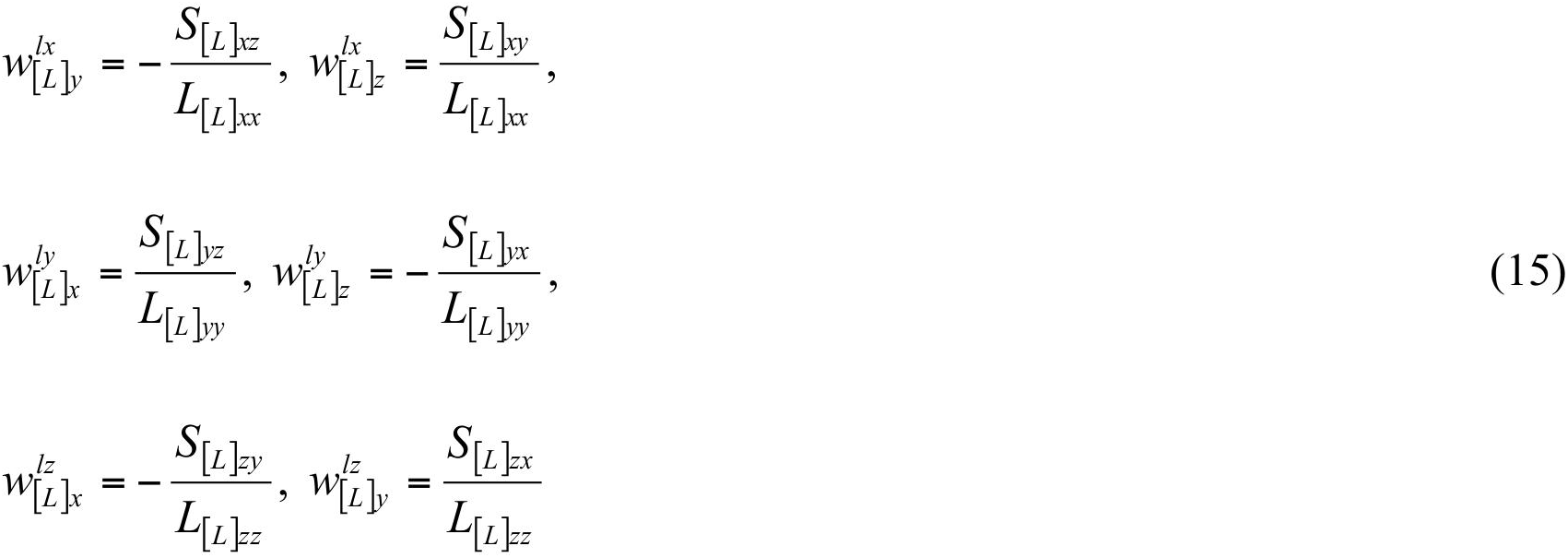

A zero value of any denominator in (15) means that there is no rotation around the corresponding axis; in this case the two corresponding numerator values must be equal to zero too, otherwise the input matrices are incompatible and the procedure must stop. If a libration axis is absent zero values are assigned to the corresponding coordinates in (15). For presentation purposes it might be useful to assign

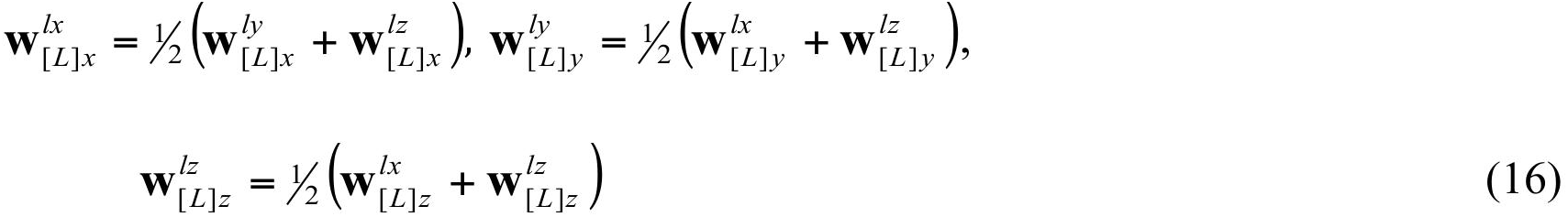

which gives the points in the middle of axes.

Knowing the position (15) of the libration axes and elements of *L*_[*L*]_ we can calculate an apparent translation due to the displacement of the libration axes from the origin

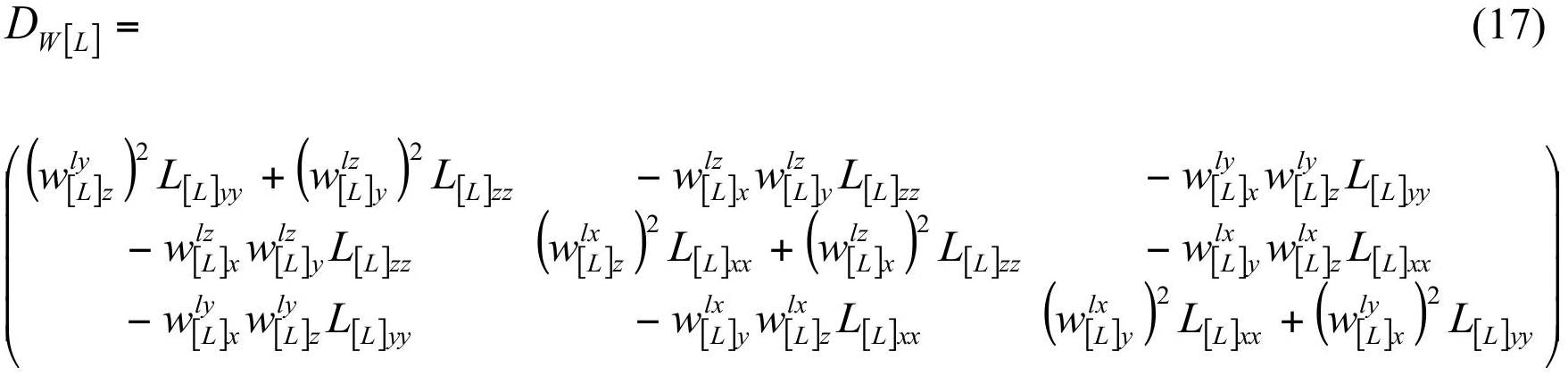

Then we can calculate the residual *T* matrix after removal of the contribution (17):

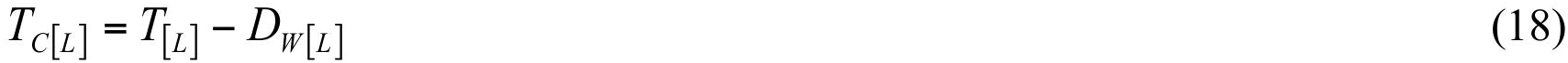

that must be positive semidefinite. Matrices *S*_[*L*]_ and *L*_[*L*]_ are not modified at this step.

## 4. Determination of the screw components (step C)

### 4.1. Correlation between libration and vibration and usual choice of S diagonal elements

Next we use the matrices *L*[*L*] and *S*[*L*] to determine the correlation parameters *s*_*x*_, *s*_*y*_, *s*_*z*_, then remove the corresponding contribution from the *T*_*C*_ matrix (equation (6)) and extract the matrix *V*_[*L*]_ for uncorrelated vibrations. However, there is an ambiguity in the definition of *S*_[*L*]_ which is apparent from the observation that the matrices *UCart*, *n* of individual atoms will not change if the same number *t* is removed simultaneously from all three diagonal elements of *S*_[*L*]_. Usually *t* is obtained by minimizing the trace of the resulting matrix *S*_*C*_ (Schomaker & Trueblood, 1968):

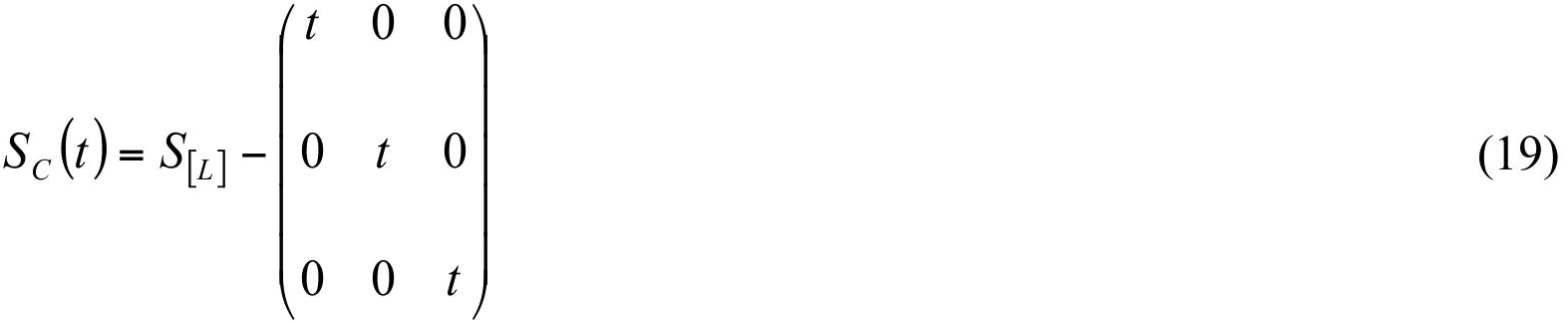

which corresponds to the minimal vibration-libration correlation (Urzhumtsev et al. 2013). This unconditioned minimization

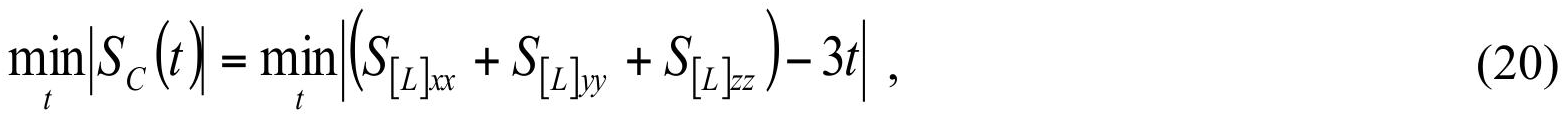

gives the value

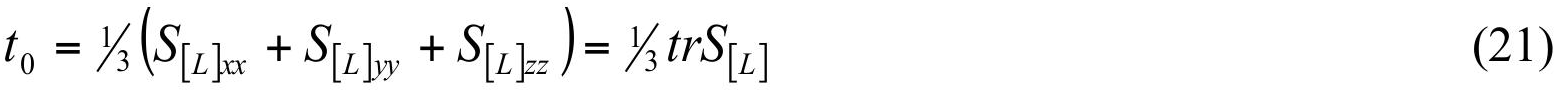

However, this value may lead to physically unrealistic matrices for which libration-vibration decomposition is impossible. First, it is clear that the *S* elements cannot be large in comparison with the elements of the *T* and *L* matrices; in particular, if there is no rotation around one of the libration axes, the corresponding *S* elements must be zero. Second, if the elements of matrix *S* and the corresponding values *s*_*x*_, *s*_*y*_, *s*_*z*_ are too large, the matrix *V* in (6) may not be positive definite. We have not identified any previous discussion of these complications in the crystallographic literature. The following sections describe a procedure that defines the constraints on the diagonal elements of matrix *S* when using (19).

### 4.2. Cauchy-Schwarz constraints

Since the diagonal elements of the positive definite matrix *V* are non-negative in any Cartesian basis, according to equations (5)–(7) the diagonal elements of the matrices must satisfy the Cauchy-Schwarz inequality

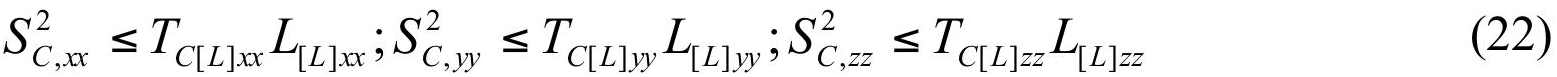

(This condition is also apparent from equations (8.5) - (8.7) in Urzhumtsev et al. 2013 which give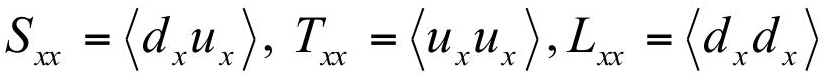 and similar conditions for other axes). In turn, this defines the limits as

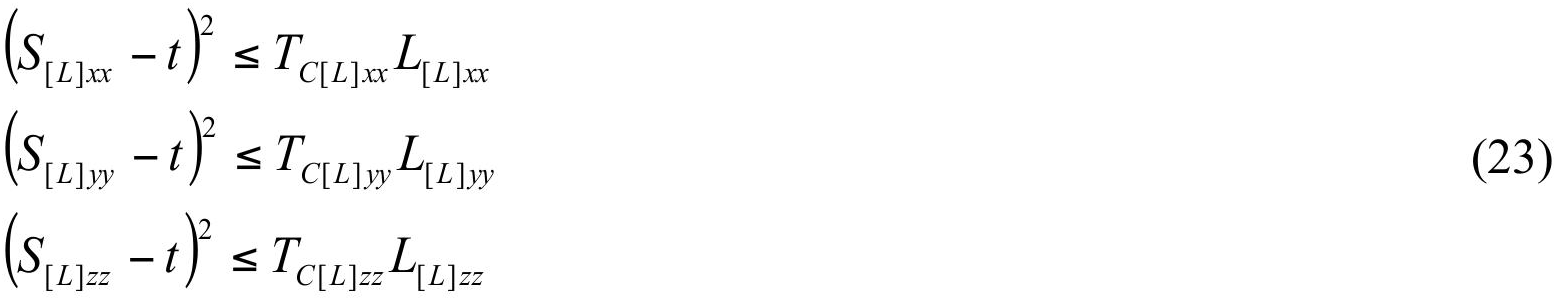

or

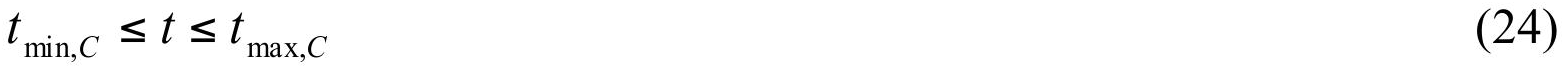

with

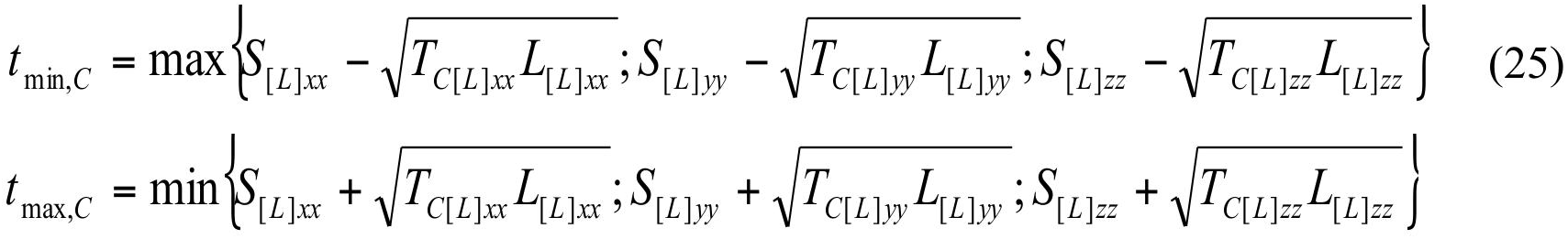

### 4.3. Positive semidefinition of the V matrix

Now we analyze the case of the matrix *V* being positive semidefinite. First, we suppose that all diagonal elements of the matrix *L*_[*L*]_ are non-zero and note that Section 4.4 considers the alternative case. From equations (5) - (7) and (19) we find the screw contribution

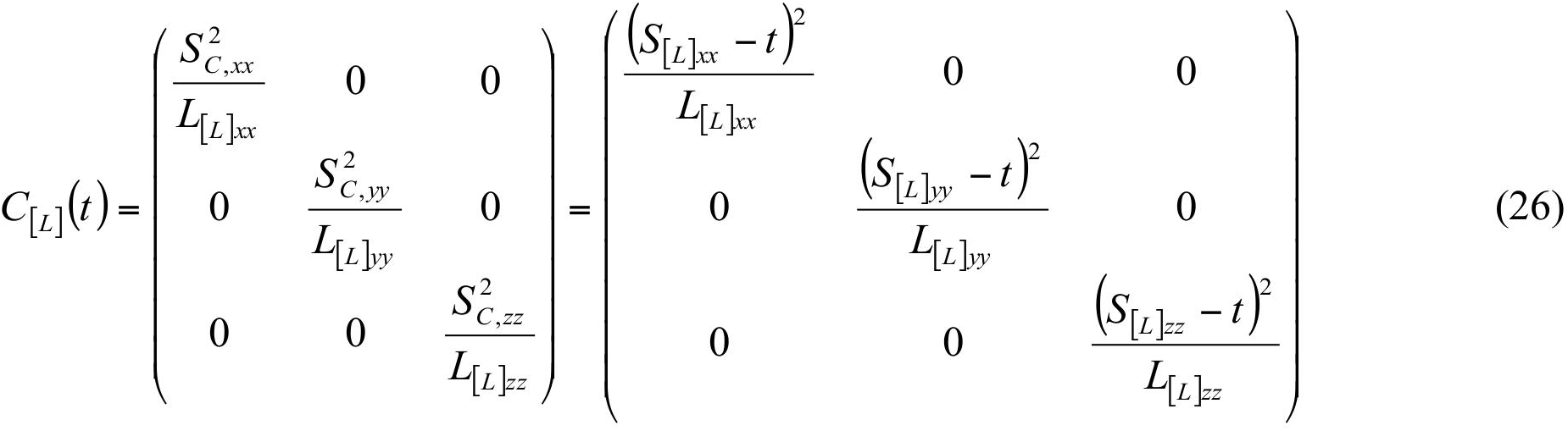

to be subtracted from matrix (18) as

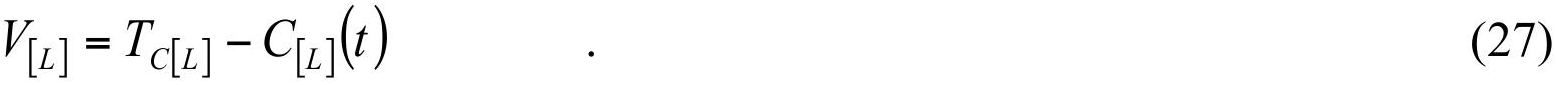

Matrix *V*_[*L*]_ is positive semidefinite along with

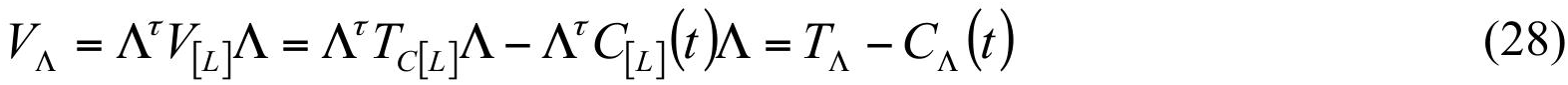

where

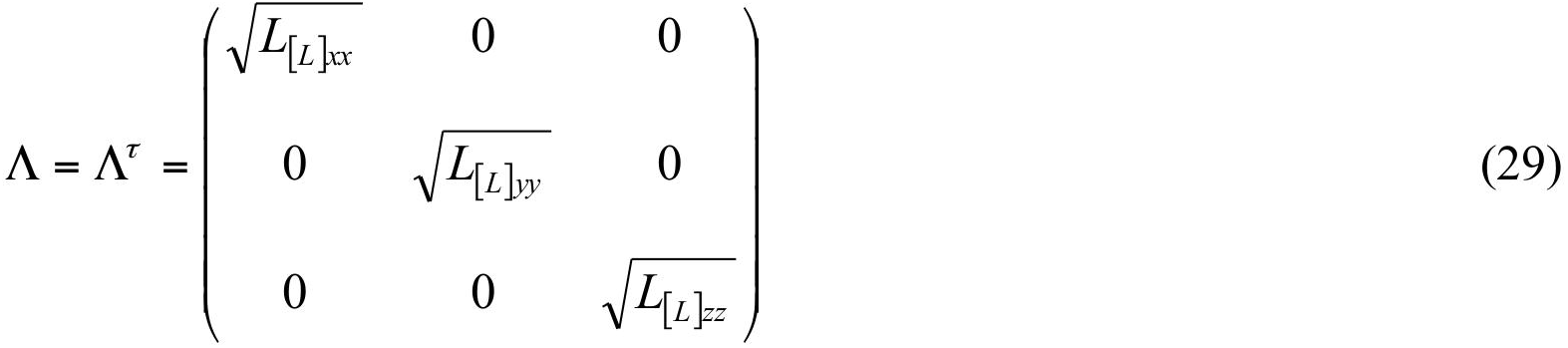

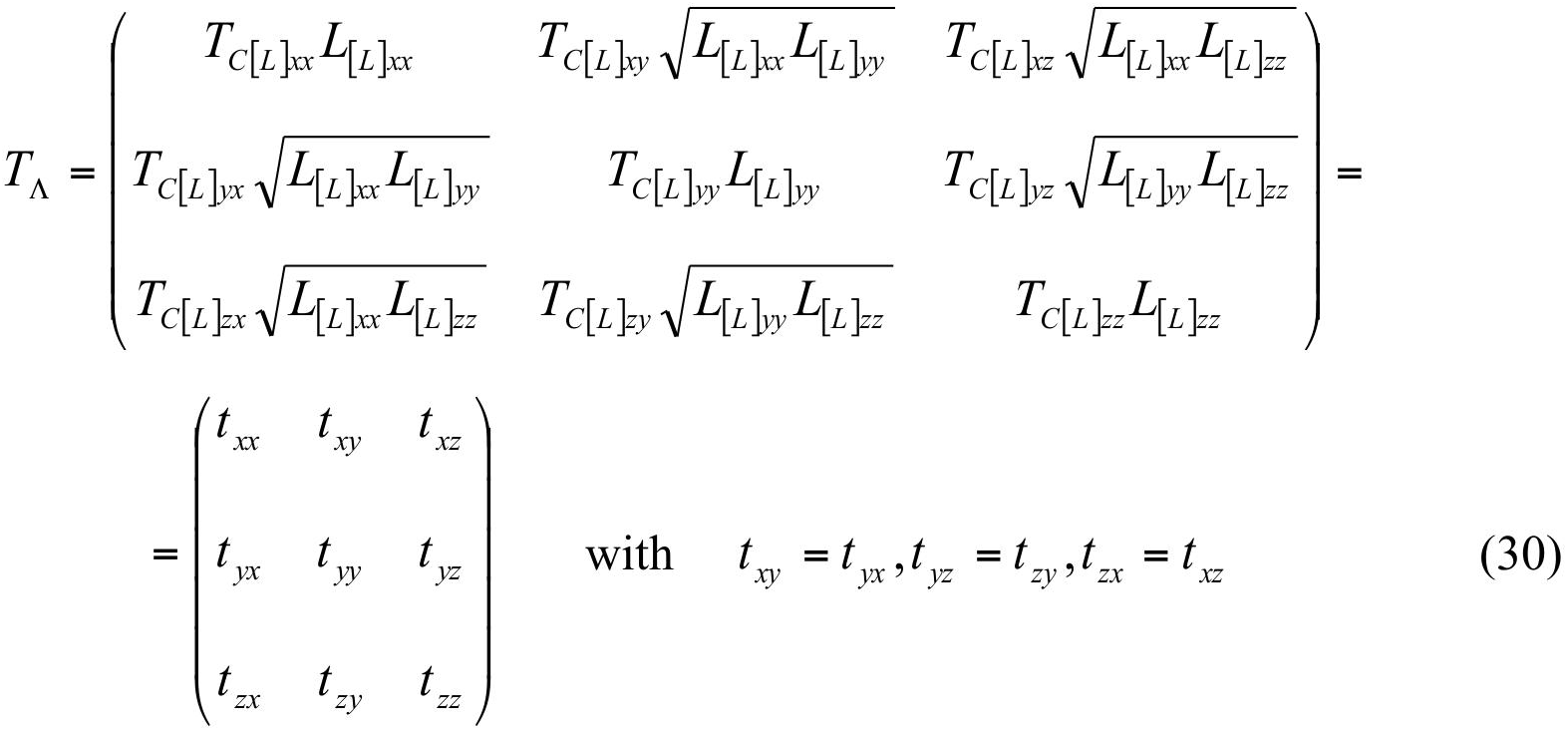

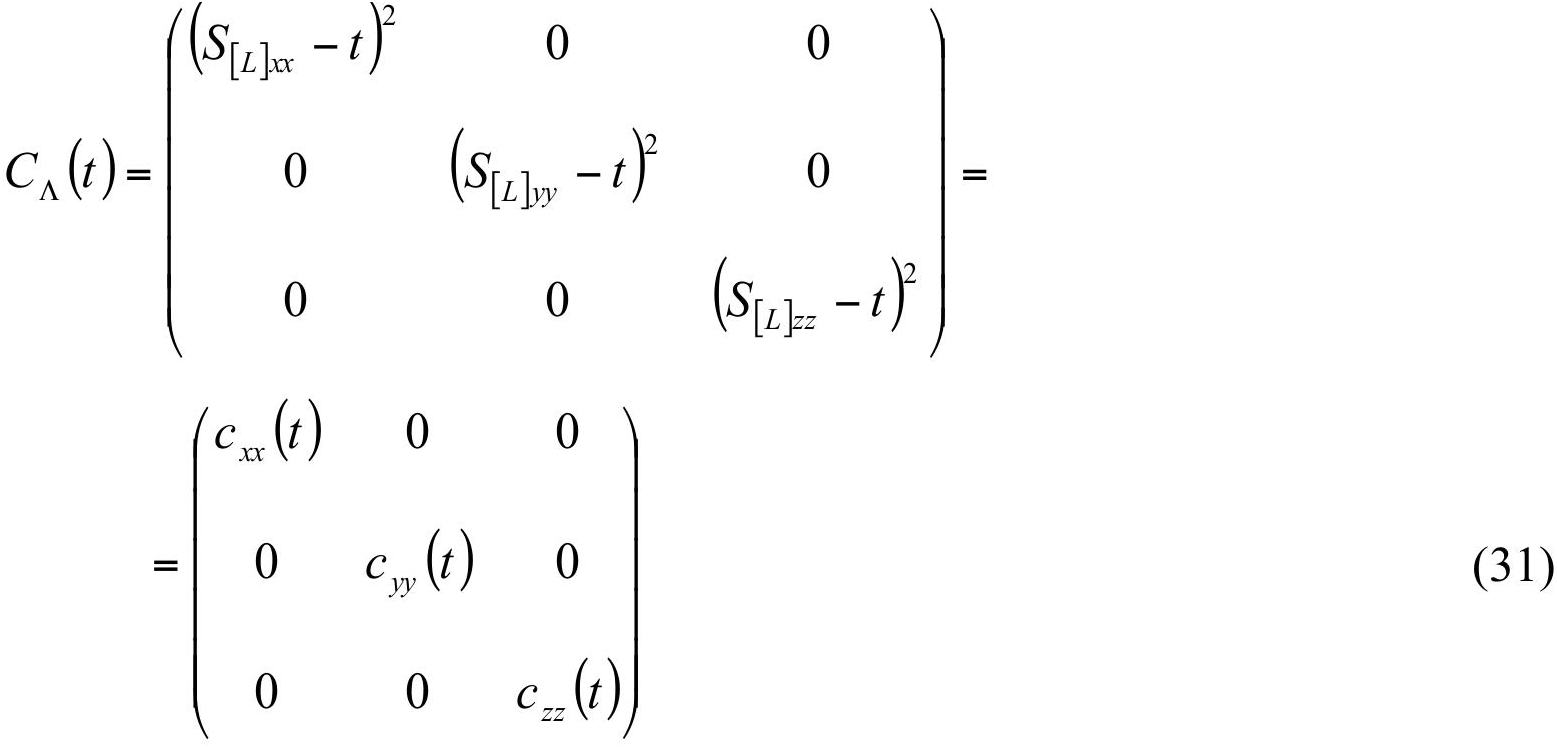

Necessarily, all diagonal terms of (31) cannot be larger than the maximal eigenvalue *τ*_max_ of (30) giving a condition

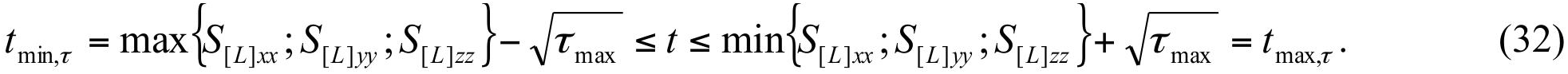

Another obvious condition that these terms are not larger than the minimum eigenvalue *τ*_min_ of (30) is sufficient but not necessary. We have not yet found a direct solution to this problem and thus use the numerical approach described below.

Matrix *V*_Λ_ is positive semidefinite if and only if all three of the real eigenvalues are nonnegative (some of them may coincide with each other). They are the roots of the cubic characteristic equation

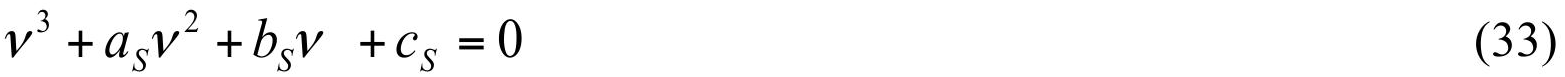

with the coefficients

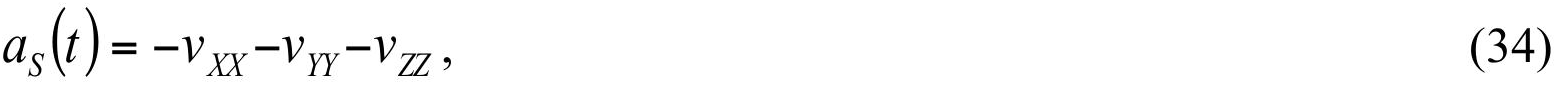

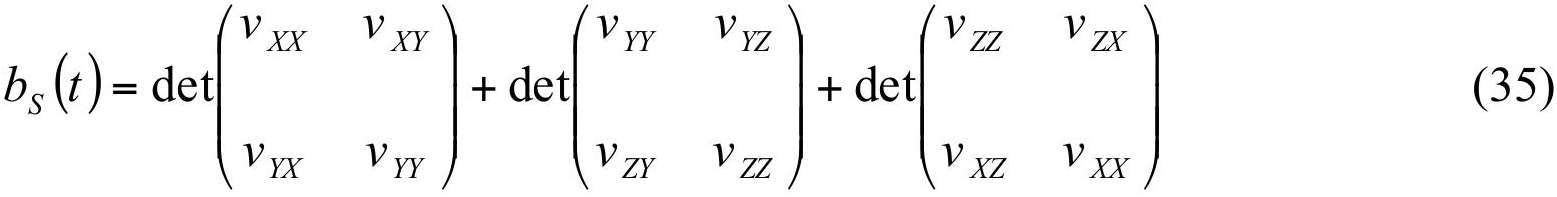

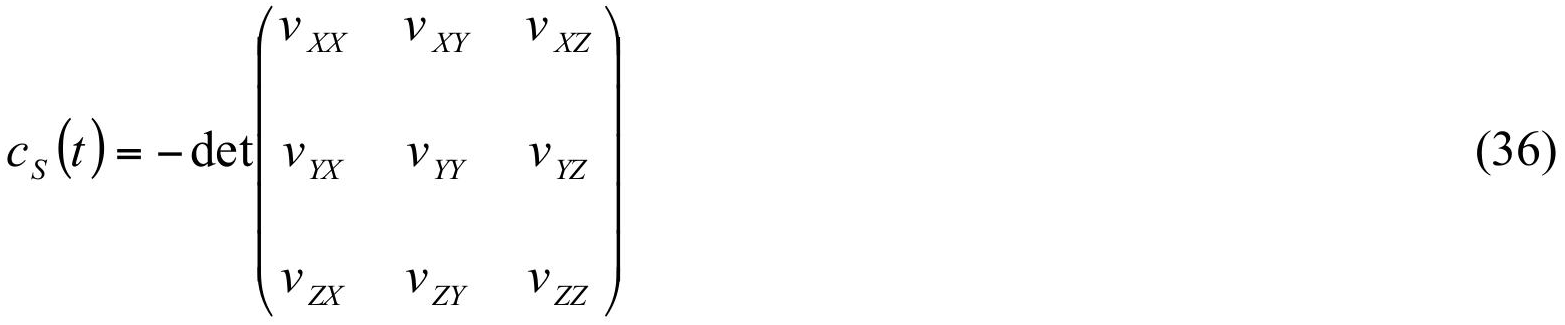

The positivity of the roots is equivalent to a system of three inequalities:

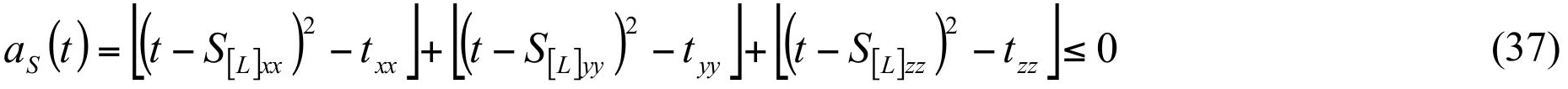

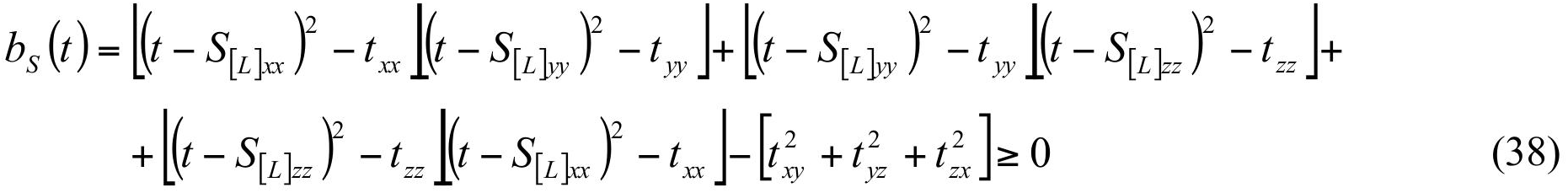

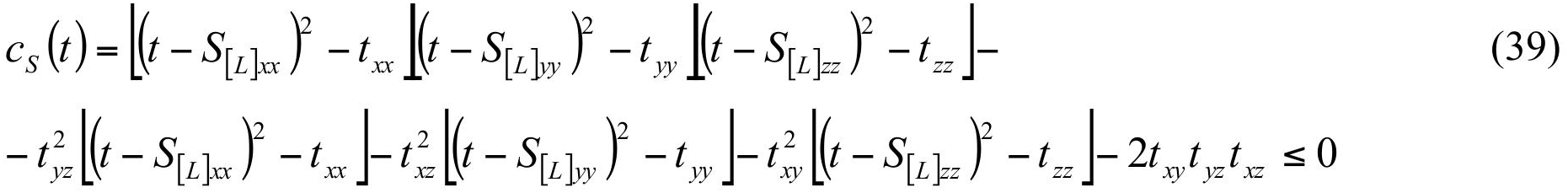

where the left parts are polynomials of order 2, 4 and 6 of the parameter *t*, all with the unit highest coefficient. The first condition (37) defines the interval for *t* values:

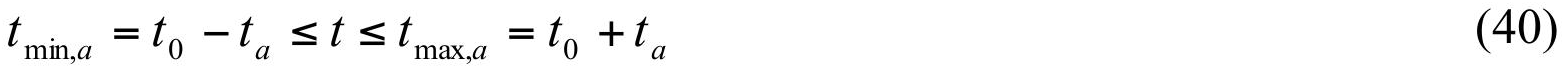

with

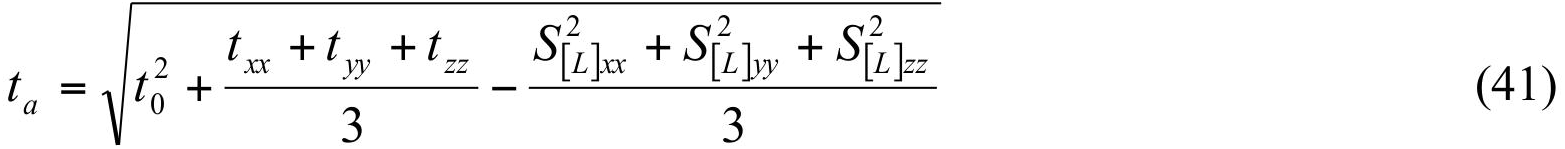

We failed to find analytical expressions corresponding to the two other inequalities and the following numerical procedure was implemented to find the best *t* value that is physically acceptable:

a) Calculate *t*_0_ value (equation (21));
b) Calculate the interval (*t*_min_,*t*_max_) for allowed *t* values as intersection of intervals (24), (32) and (40); *t*_min_ = max{*t*_min,*C*_,*t*_min,*τ*_,*t*_min,*a*_ }, *t*_max_ = min{*t*_max,*C*_, *t*_max,*τ*_, *t*_max,*a*_ }; if *t*_min_ > *t*_max_ the problem has no solution and the procedure stops;
c) If *t*_min_ = *t*_max_ we check the conditions *b*_*S*_ (*t*_min_) ≥ 0, *c*_*S*_ (*t*_min_) ≤ 0, or directly the condition that *V*_Λ_ is positive semidefinite; if the conditions are satisfied we assign *t*_*S*_ = *t*_min_ otherwise the problem has no solution and the procedure stops;
d) If *t*_min_ < *t*_max_ we search numerically, in a fine grid, for the point *t*_*S*_ in the interval (*t*_min_,*t*_max_) and closest to *t*_0_ such that *b*_*S*_ (*t*_*S*_) ≥ 0, *c*_*S*_ (*t*_*S*_) ≤ 0; if at any point of this interval at least one of these inequalities is wrong, then the procedure stops;
e) If steps *c* and *d* give us the *t*_*S*_ value, we accept it as the solution of the problem otherwise there is no acceptable *t* value and the procedure of the *TLS* matrices decomposition stops.

### 4.4. Singular sets of rotation

When one of the *L*_[*L*]*xx*_, *L*_[*L*]*yy*_, *L*_[*L*]*zz*_ values is zero (that is, there is no rotation around the corresponding axis), straightforward use of the standard procedure including (26) becomes impossible. However, in this case the *t*_*S*_ value must equal to *S*_*C*,*xx*_, *S*_*C*,*yy*_ or *S*_*C*,*zz*_, corresponding to the axes with no rotation, turning the corresponding inequality in (25) into an equality and making the corresponding diagonal element in (26) equal to zero; for example if *L*_[*L*]*xx*_ = 0 then *t*_*S*_ = *S*_*L*,*xx*_ resulting in *S*_*C*,*xx*_ = 0. We simply need to check two other conditions in (22) and the condition that the residual matrix is positive semidefinite, for example calculating (37)-(39). If *t*_*S*_ does not satisfy these conditions, the problem has no solution and the procedure of the *TLS* matrices decomposition stops.

### 4.5. Screw parameters

For the *t*_*S*_ determined above we calculate the matrix *S*_*C*_ (*t*_*S*_) (19). From this matrix we obtain the estimates 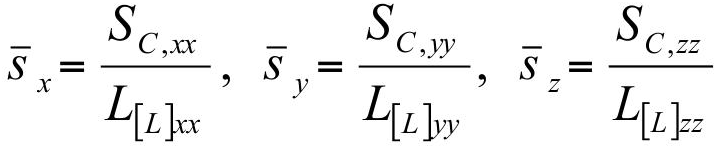 for the screw parameters following the rotation axes currently aligned with the coordinate axes of the basis *L*. We remind the reader that if one of the *L*_[*L*]*xx*_, *L*_[*L*]*yy*_^,^*L*_[*L*]*zz*_ values is equal to zero, the corresponding diagonal element of *S*_*C*_ must also be equal to zero (otherwise the matrices are inconsistent with each other and the procedure stops) and we assign the corresponding screw parameter, 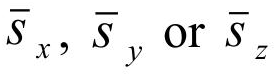, to be zero.

## 5. Determination of the vibration components (step D)

### 5.1. Matrix V and vibration parameters in L basis

When the *t*_*S*_ value is known, matrix *C*_[*L*]_ (*t*_*S*_) and then *V*_[*L*]_ are calculated according to equations (26) - (27). The step of obtaining independent vibrations from the *V*_[*L*]_ matrix is similar to that for getting the independent librations from *L*_[*L*]_. First, we calculate the three eigenvalues 0 ≤ *m*_1_ ≤ *m*_2_ ≤ *m*_3_ of matrix *V*_[*L*]_ (in practice, all of them are strictly positive). Then we identify three corresponding unit eigenvectors **v**_1_, **v**_2_, **v**_3_ that are orthogonal to each other and assign

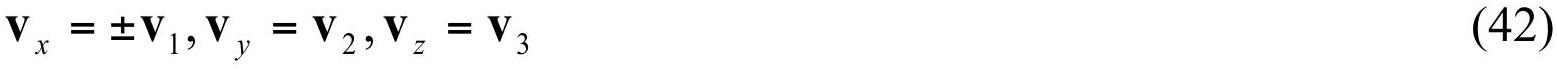

(the sign for **v**_**x**_ is taken so that the vectors (42) form a right-hand triad). The three eigenvectors give the directions of the uncorrelated translations and the corresponding eigenvalues *m*_1_, *m*_2_, *m*_3_ are squared *rmsd*s along these axes. These axes define a new orthonormal basis V in which matrix *V*_[*L*]_ becomes the diagonal matrix *V*_[*V*]_ with elements *V*_[*V*]*xx*_ = *μ*_1_*, V*_[*V*]*yy*_ = _2_,[*V* ]*zz* = *μ*_3_; these values are the estimates 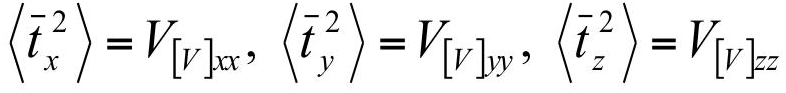 of the squared vibration amplitudes.

### 5.2. Vibration and libration axes in M basis

The libration and vibration amplitudes and screw parameters are independent of the choice of the basis, and the direction of the libration axes (11) is known in the principal M basis. However the directions of the uncorrelated translations **v**_1_, **v**_2_, **v**_3_ that were calculated in section 4 and the points 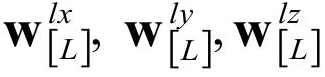 belonging to the libration axes (section 3.2) are given in the L basis.

To obtain the coordinates 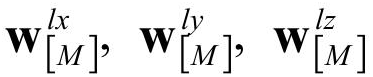 of these points in the M basis we apply the transformation

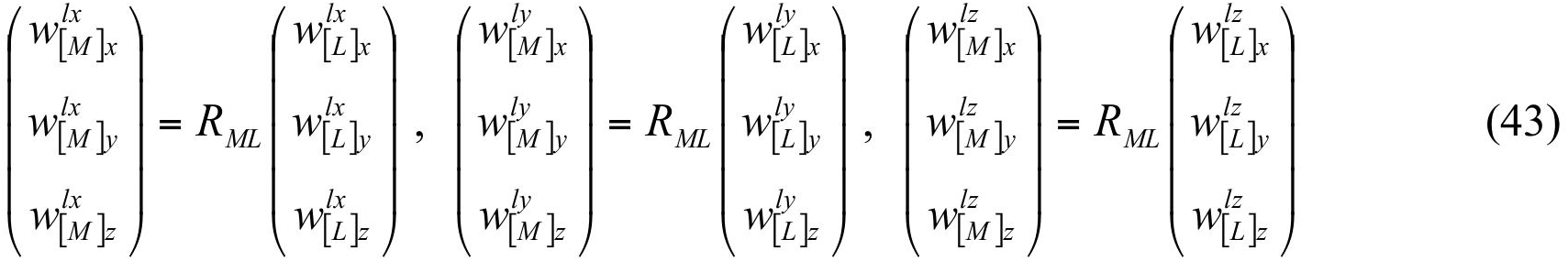

using (12). Similarly, the direction of the axes **v**_*x*_, **v**_*y*_, **v**_*z*_ in the M basis and can be obtained as

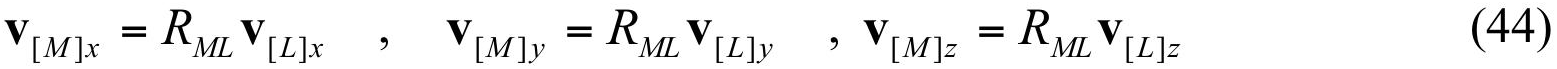

### 5.3. Calculations in V basis

Knowledge of **v**_*x*_, **v**_*y*_, **v**_*z*_ (42) in the basis M defines the rotation matrix from the M basis to the V basis as

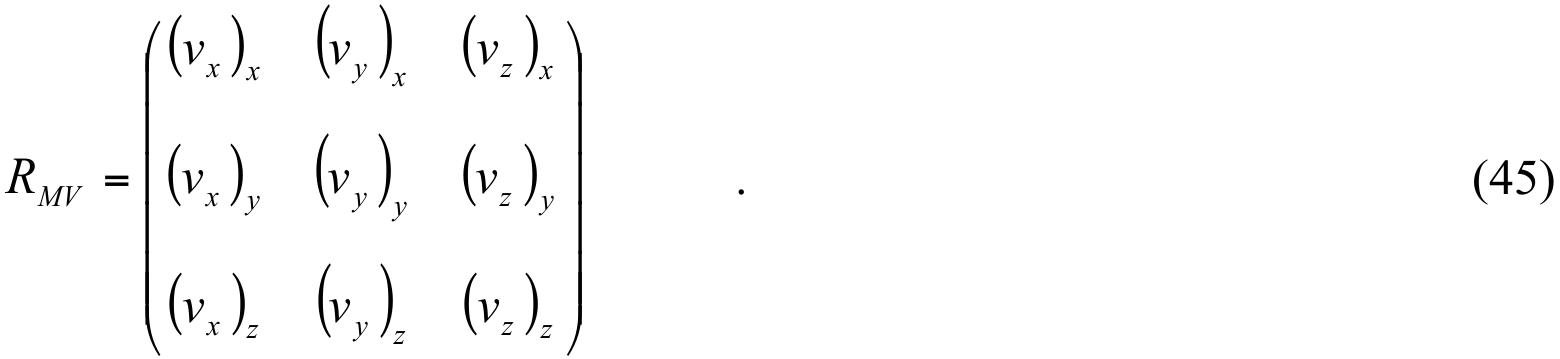

The coordinates of a vector expressed in the two bases are linked by the equation

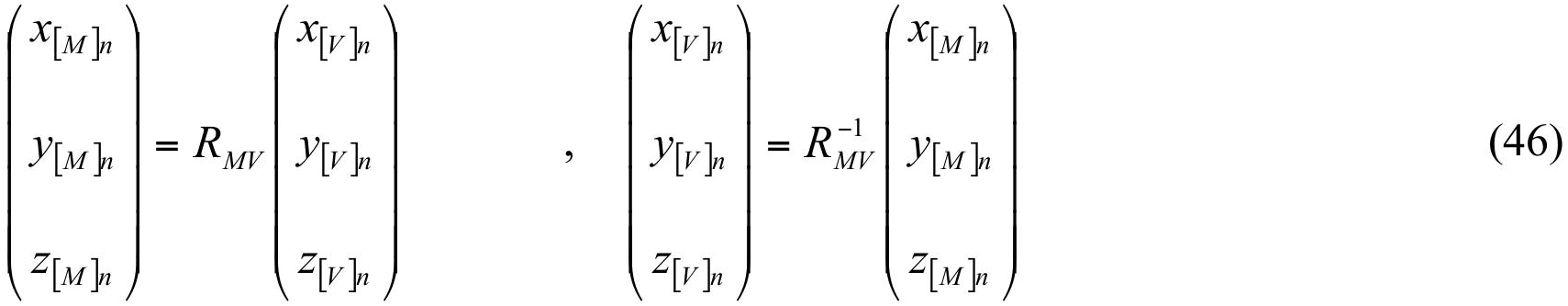

This step finalizes extracting the parameters of the motions that correspond to the given set of *TLS* matrices. Section 6 provides some examples of this procedure’s application to models deposited in the PDB.

## 6. Examples of *TLS* matrix decomposition

### 6.1. General procedure

To illustrate the algorithm described above we analyzed several entries selected from the PDB. For each structure, we applied a standard *TLS* refinement protocol from *Phenix* (Adams et al.,2010; Afonine et al., 2012) including automatic determination of the *TLS* groups. During refinement, 20 matrix elements (6 for *T*, 6 for *L* and 8 for *S* where the third diagonal element of *S* has been chosen to make the trace of the matrix equal to 0) were refined independently. The procedure described above was then applied to all sets of the *TLS* matrices obtained. By current conventions, the *L* and *S* matrices for the PDB models are expressed in degrees^2^ and in Å·degree, respectively, and first we need to convert them into radians^2^ and Å·radians.

### 6.2. Synaptotagmin

The crystals of synaptotagmin III (PDB code 1dqv) contain two copies of the molecule in the asymmetric unit. The structure, when re-refined by *phenix.refine*, has *R*_*work*_ = 0.200 and *R*_*work*_ = 0.231. The *TLS* refinement, with each molecule taken as a single *TLS* group, reduced *R*-factors to *R*_*work*_ = 0.177 and *R*_*work*_ = 0.211 and shows the significance of this *TLS* modeling. Table 2 shows the two sets of the matrices, and Table 3 shows the corresponding motion parameters extracted using our routine. For the two groups both vibrations and librations are practically isotropic and are of the same order of magnitude for both groups. Fig. 2a shows the principal axes of these motions.

**Figure 2.**
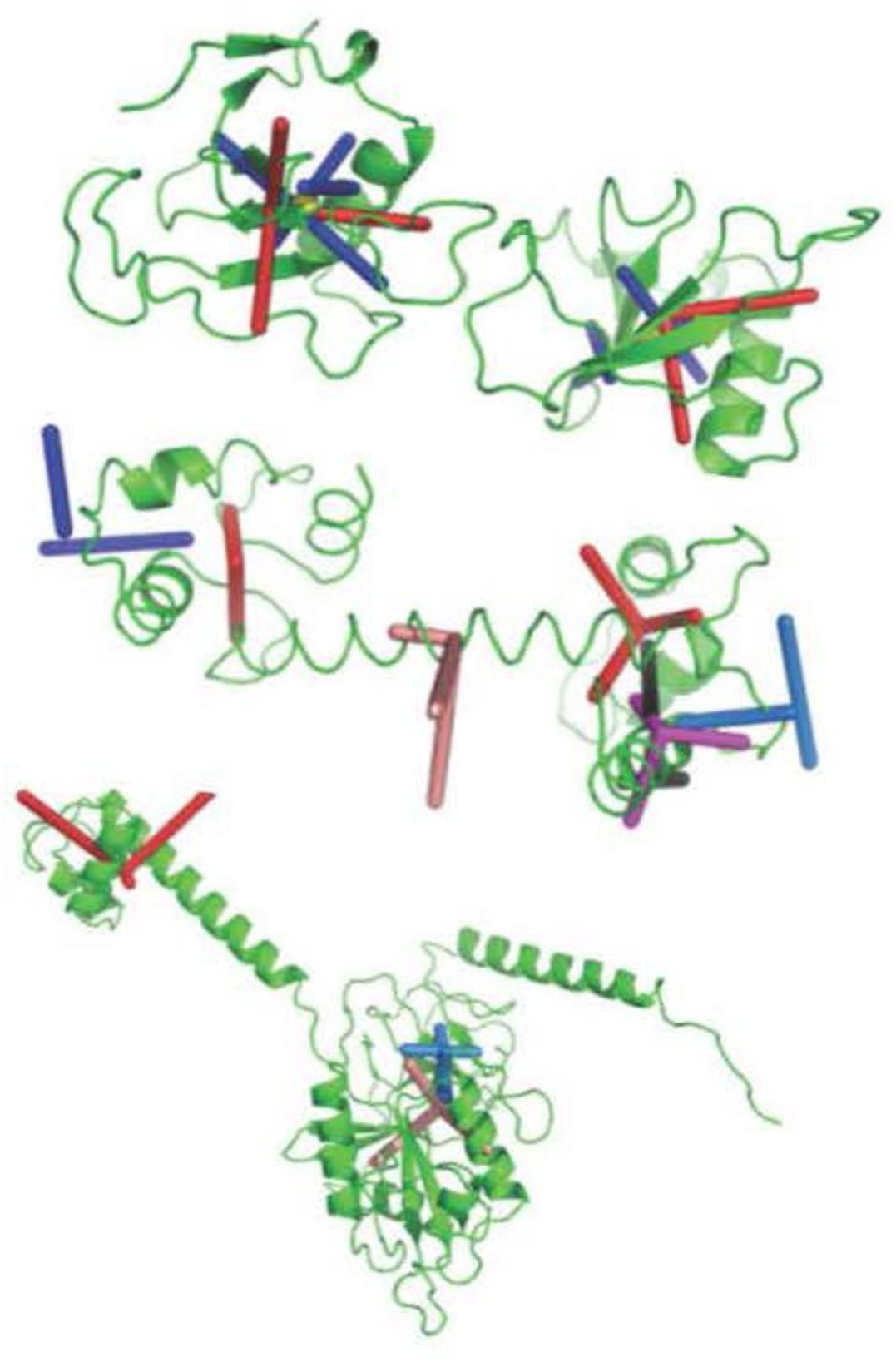
Examples of the vibration-libration ensembles. Red / salmon / magenta sticks indicate the principal vibration axes with the origin in the centre of the group; blue / marine / black sticks are for the libration axes. Yellow spheres for the 1dqv model show the reaction centers. a) 1dqv model. (b) 1exr model; note pure vibrations for the group 3 (the helix) and absence of one of libration axes for the groups 1 and 2. c) 4b3x model. Libration axes for the first group are not shown being too far from the molecule.

**Table 2.** Examples of the *TLS* matrices.

| PDB<br>code | chain,<br>residue<br>number | $T(\text{\AA}^2)$ | $L(\text{degree}^2)$ | $S(\text{\AA}\cdot\text{degree})$ |
| --- | --- | --- | --- | --- |
| 1dqv | A1-A97 | 0.1777 0.0090 -0.0044<br>0.0090 0.1306 0.0019<br>-0.0044 0.0019 0.1372 | 1.4462 -0.0160 -0.2656<br>-0.0160 1.2556 0.4713<br>-0.2656 0.4713 0.8689 | 0.0467 -0.0523 0.0566<br>0.1010 0.0032 -0.0164<br>0.0090 0.0188 0.0560 |
|  | B1-B97 | 0.1777 0.0090 -0.0044<br>0.0090 0.1306 0.0019<br>-0.0044 0.0019 0.1372 | 1.4462 -0.0160 -0.2656<br>-0.0160 1.2556 0.4713<br>-0.2656 0.4713 0.8689 | 0.0467 -0.0523 0.0566<br>0.1010 0.0032 -0.0164<br>0.0090 0.0188 0.0560 |
| 4b3x | A1-A65 | 0.4663 0.0991 -0.0764<br>0.0991 0.5443 -0.0321<br>-0.0764 -0.0321 0.5001 | 0.4738 0.0063 0.2318<br>0.0063 0.2120 -0.0584<br>0.2318 -0.0584 0.1312 | 0.0391 -0.0307 -0.4316<br>0.0587 0.1786 -0.2003<br>0.3665 0.4293 0.0403 |
|  | A66-A363 | 0.1649 -0.0259 0.0184<br>-0.0259 0.1422 0.0055<br>0.0184 0.0055 0.2028 | 0.8808 -0.0912 -0.1736<br>-0.0912 0.9522 0.0972<br>-0.1736 0.0972 1.6563 | -0.0345 0.0102 -0.0661<br>0.1159 -0.0222 0.0999<br>0.0424 -0.1330 -0.0237 |

**Table 3.**
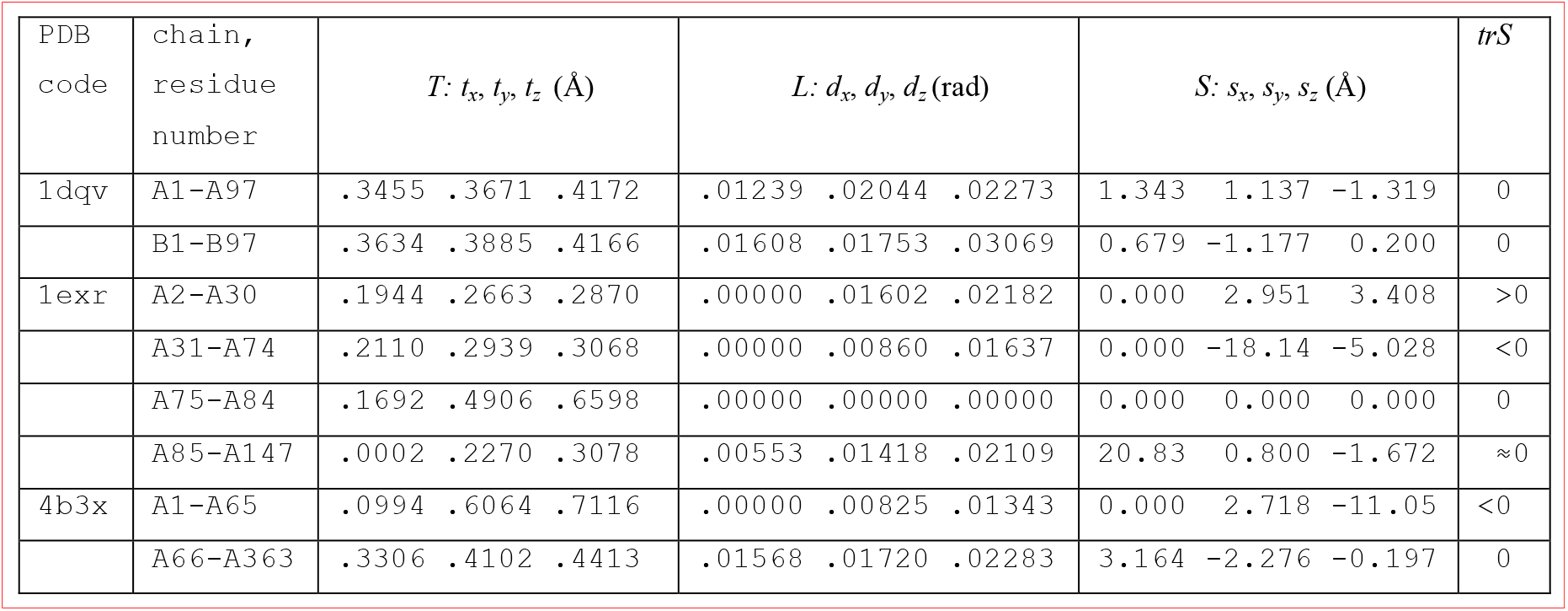
Examples of parameters of the elementary motions found form the decomposition of the *TLS* matrices.

### 6.3. Calmodulin

The structure of calmodulin (PDB code 1exr) was automatically split into 4 *TLS* groups. This case was an example of possible problems that could be solved by a minimal manual intervention. For the first group, one of the eigenvalues of the matrix *L* was marginally negative, equal to - 0.000023; we allowed the program to substitute the value 0 thus accepting this group to be singular with one degenerate libration. The second group also had one degenerate libration. After correcting the slightly negative eigenvalue for the first group, all composited motions were extracted without problems.

For the third group, the screw parameters and axes positions were extremely large, leading to the procedure’s inability to find a positive semidefinite *V*_[*L*]_. This was caused by the fact that all three eigenvalues of the matrix *L* were practically equal to 0 (0.000000, 0.000008 and 0.000035) resulting in high computational instability. We replaced matrix *L*, and respectively *S*, by zero matrices, defining all librations as absent, and obtained significant values for vibration parameters. This group is a helix held at both ends by large domains, which explains the impossibility of librations within the helix.

Finally, for the fourth group one of the diagonal elements of the matrix *T* was marginally negative. Increase all diagonal elements of the matrix *T* by 0.002 makes this matrix positive definite. This manipulation can be considered a form of ‘borrowing’ roughly 0.16 Å^2^ from either the overall scale factor or from individual atoms (*i.e.* such that the *T* modification and the simultaneous removing of 0.16 Å^2^ from all corresponding ADP parameters does not change the structure factors). An accurate separation of total ADP values into contribution from several sources (for example, Winn et al., 2001, 2003; Afonine et al., 2013) is a separate ongoing project (Afonine & Urzhumtsev, 2007). This group vibrates essentially in a plane (Fig. 2b) and the principal vibration axis of the group 3 (the helix) is parallel to this plane, leading to the plausible hypothesis that these groups at least partially move together.

### 6.4. Initiation translation factor 2 (IF2)

The structure of IF2 (PDB code 4b3x) has recently been solved in one of our laboratories (Simonetti *et al*., 2013) with *R*_*work*_ = 0.180 and *R*_*free*_= 0.219 at a resolution of 1.95 Å. *TLS* refinement was performed with two groups: the first included the N-terminal and the following long helix, and the second included the rest of structure. This refined model had lower *R*_*work*_ = 0.176 and *R*_*free*_= 0.203. For this example, the *TLS* matrices from the first group were not directly interpretable because the residual matrix *V*_[*L*]_ was not positive semidefinite having one of its eigenvalues equal to −0.05. Similarly to the last group of the calmodulin, we artificially added 0.06 to all diagonal elements of the matrix *T* which corresponds to ‘borrowing’ of about 5 Å^2^ from individual atoms or from the overall scale factor (as above). With such a correction the procedure worked well. It should be noted that for the first *TLS* group one of the rotations was degenerated and that the assignment *trS* = 0 would make this matrix incompatible with *L*. Table 3 shows that vibrations of this group are essentially anisotropic. Fig. 2c also shows that the libration axes for this group pass quite far away from the molecule, which makes the corresponding rotation similar to a translation. Additionally, the large 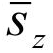 value leads us to believe that the matrix *S* is not well defined to be physically significant. The matrices for the second group were routinely interpreted showing isotropic vibrations and librations.

We then tried to apply the same procedure after manually choosing the *TLS* groups as residues 1-50 (N terminal), 51-69 (helix), 70-333 (G domain) and 343-363 (connector to the C domain absent in this structure). As before, the G domain the matrices were physically interpretable. For the other two groups, after a manual adjustment similar to those discussed above (a slight artificial increasing of the diagonal *T* elements with an accompanying decrease in the residual ADP of the individual atoms), we obtained a pure vibration for the helix (as for the calmodulin case) and a libration around a single axis for the terminal group. In contrast, we failed to find reasonably small corrections for the matrices of the first group that would make them physically interpretable.

## 7. Interpretation of *TLS* matrices with an ensemble of models

### 7.1. Generation of an explicit set of atomic models with a given variability consistent with TLS

Some structural problems may explicitly require a set of models that describe a given mobility, *e.g.* corresponding to the *TLS* matrices for harmonic motion. We do not discuss here larger-scale anharmonic motions for which other techniques such as molecular dynamic trajectories have traditionally been used, as originated by McCammon et al., 1977. An example of such a problem is described in the accompanying paper by Van Benschoten *et al*. in which X-ray diffuse scattering data were compared with calculated data corresponding to different types of molecular motion.

As soon as vibration and libration parameters are known, we can build a corresponding set of models explicitly. If a model deposited in the PDB contains *TLS* matrices, the procedure of their decomposition described above can be applied. A decomposition of this motion into three independent vibrations and three independent librations provides the corresponding atomic shifts, the sum of which gives the total displacement.

It is generally more convenient to generate each group of atomic shifts in its own basis: shifts 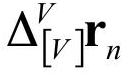 due to vibration in the V basis and shifts 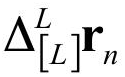 due to libration in the L basis. Here we are working in a linear approximation such that rotation angles are of order of 0.1 radian. For each particular set of generated shifts, they are the transformed into the M basis as 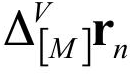 and 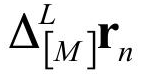 and their sum

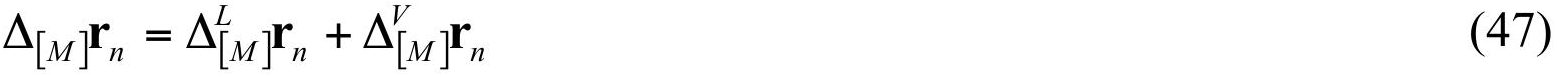

is applied to the corresponding atoms. Details of the generation of such a model are discussed in the next sections. This procedure is repeated independently many times generating randomly moved models distributed accordingly to the *TLS* matrices.

### 7.2. Calculation of the model shift due to libration

Let us suppose that, using the procedure described above, we know the three mutually orthogonal axes ***l***_**x**_, ***l***_**y**_, ***l***_**z**_ for independent libration and the points that belong to these axes. If the coordinates of these points 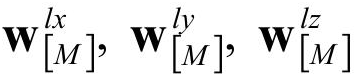 are given in the molecular basis M we can recalculate them into the L basis 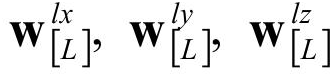 using (14). We can do the same for the coordinates of all atoms of the group that are subject to the *TLS* motion: (*x*_[*L*]*n*_, *y*_[*L*]*n*_, *z*_[*L*]*n*_), *n* = 1, 2, …, *N*. We note that the squared libration amplitudes 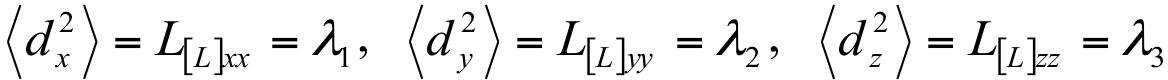 (Section 3.2) and the screw parameters *s*_*x*_, *s*_*y*_, *s*_*z*_ (estimates for which 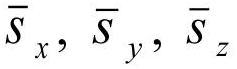 are defined in Section 4.5) are independent of the chosen basis.

For an atom at a distance *R* = 1 Å from the rotation axis, the probability of the shifts *d*_*x*_*, d*_*y*_*, d*_*z*_, which are numerically equal to the rotation angle in radians, are equal to:

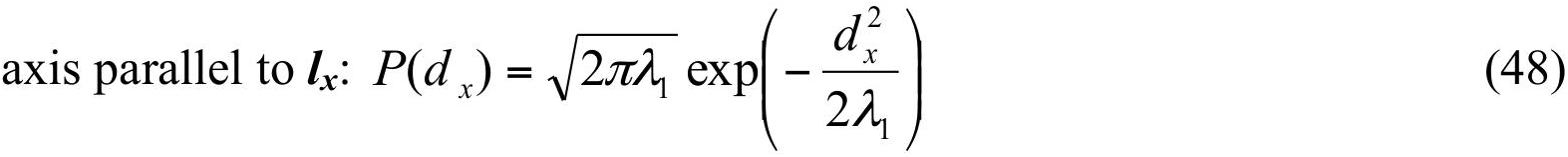

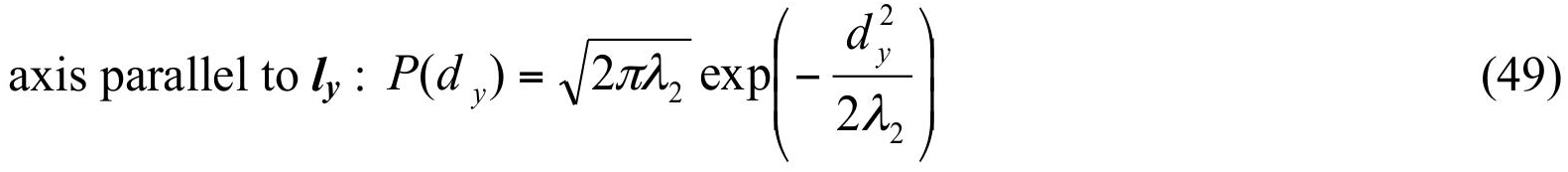

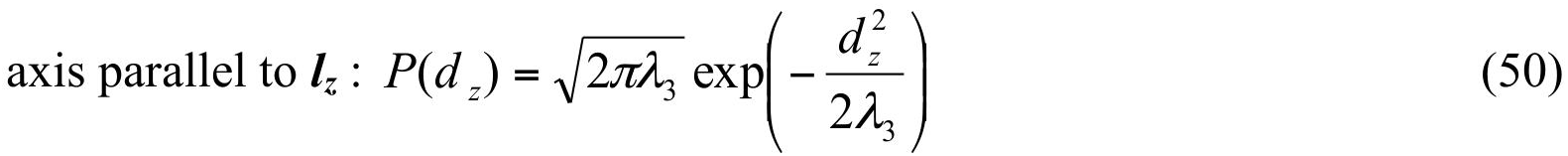

If one of the eigenvalues is equal to 0 then the corresponding *d* is equal to 0 with unit probability. Let us obtain particular values of *d*_*x*0_, *d*_*y*0_, *d*_*z*0_ using a random numbers generator with the normal distribution (48-50).

For the rotation around the axis parallel to ***lz*** and crossing the point 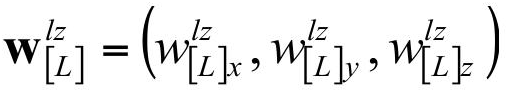, for each atom *n* described by the vector **r***n* we recalculate the coordinates of 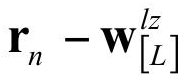 in the L basis

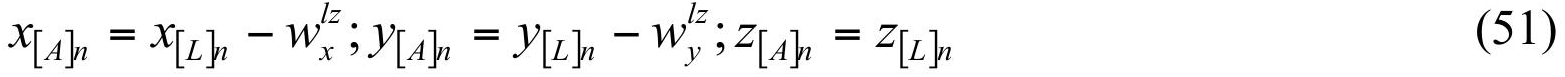

If 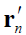 stands for the position of the same atom after rotation by angle *d*_*z*0_ around this axis, the coordinates of 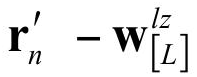 can be written as

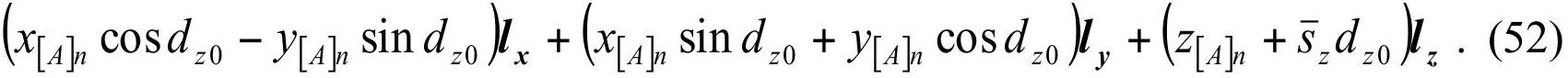

This gives the corresponding atomic shift 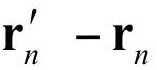

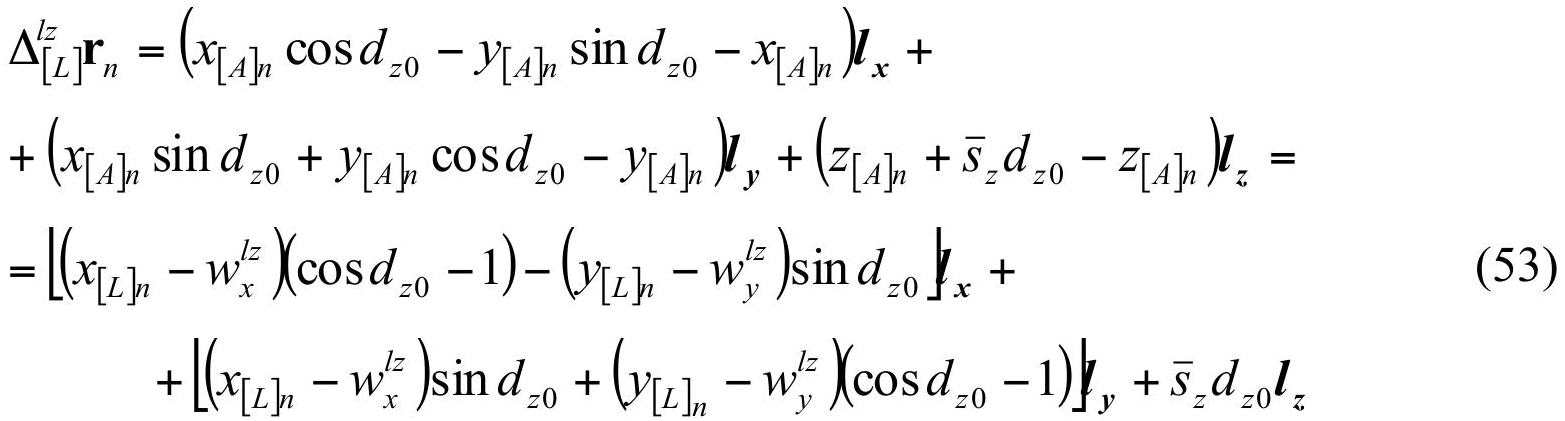

Similarly for two other axes:

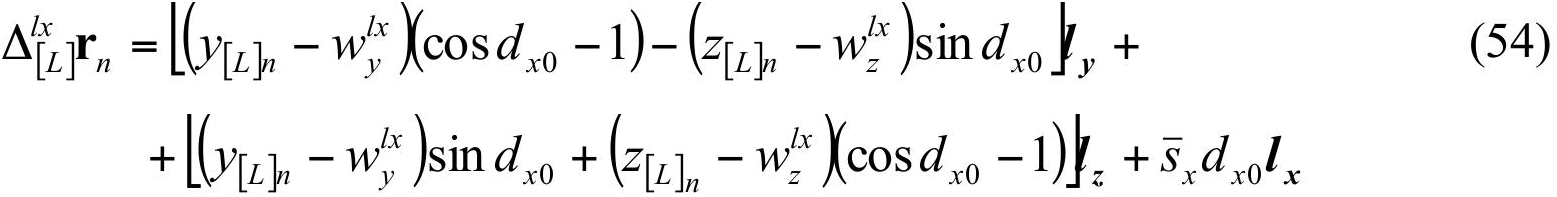

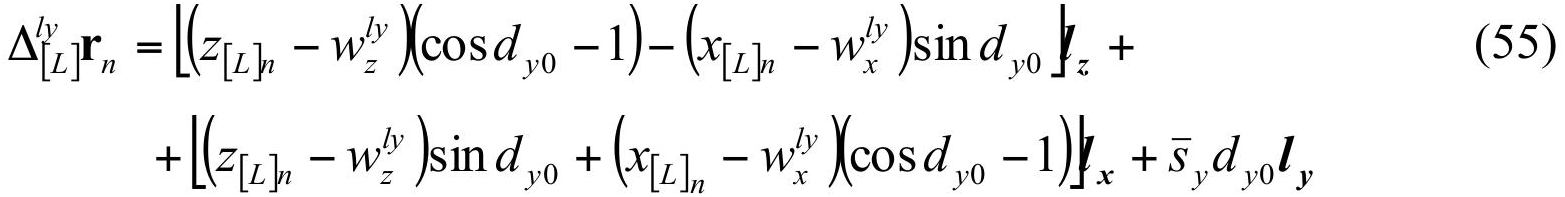

The overall shift due to libration (around the three axes) is the sum of shifts (53)-(55)

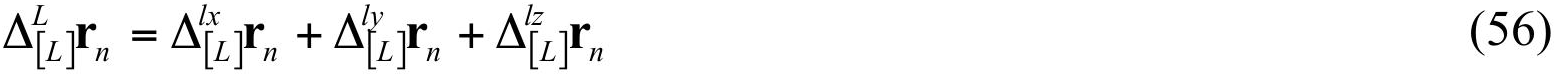

It changes from one atom of the group to another and must be calculated for all atoms of the group with the same (*d*_*x*0_, *d*_*y*0_, *d*_*z*0_) values for a particular instance of the three rotations.

The atomic shift (56) is given in the L basis. To transform this shift from the L basis into the initial M basis, we invert equation (14):

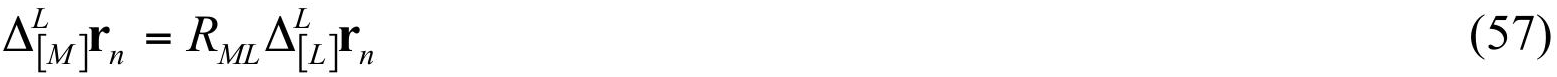

### 7.3. Calculation of the model shift due to vibration

In the harmonic approximation, the independent vibration shifts *t*_*x*_, *t*_*y*_, *t*_*z*_ expressed in the V basis are distributed accordingly to the probability laws:

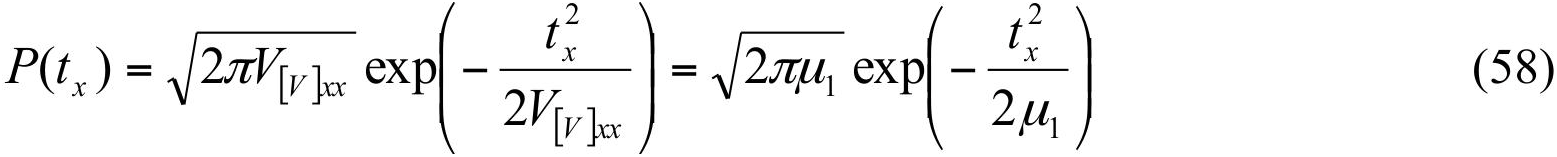

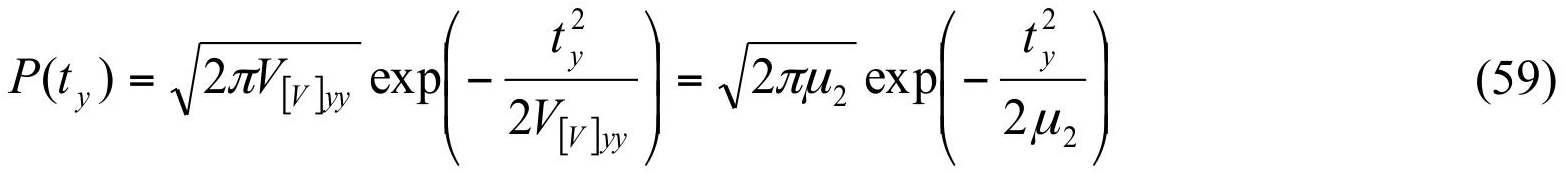

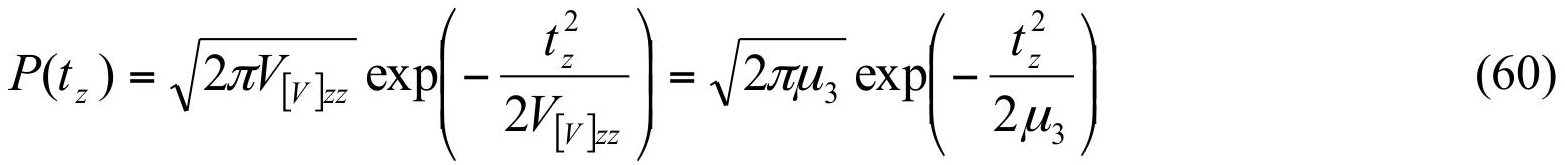

Using a random numbers generator, for each model we obtain particular values of *t*_*x*0_, *t*_*y*0_, *t*_*z*0_ using (58-60). If one of the eigenvalues *μ* is equal to zero, the zero value is assigned to the corresponding shift. The overall translational shift, common to all points of the rigid group, is equal to

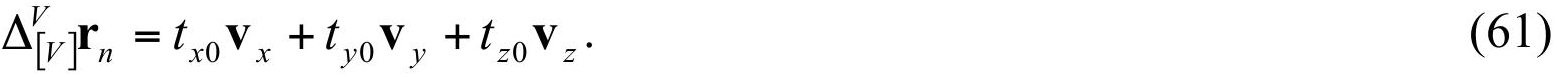

In order to obtain this shift in the M basis we calculate, similarly to equation (57),

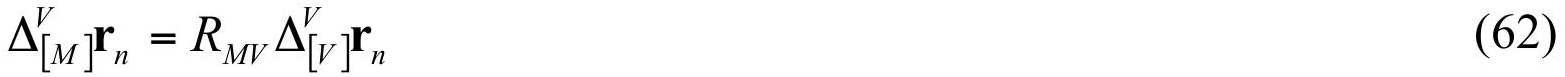

### 7.4. Validation and application to GpdQ

We generated the ensembles produced by alternative *TLS* refinements of the glycerophosphodiesterase GpdQ (Jackson et al., 2007). GpdQ is found in *Enterobacter aerogenes* and contributes to the homeostasis of the cell membrane by hydrolyzing the 3’-5’ phosphodiester bond in glycerophosphodiesters. Each dimer contains three distinct domains per monomer: an α/β sandwich fold containing the active site, a domain-swapped active site cap and a novel dimerization domain comprised of dual-stranded antiparallel β-sheets connected by a small β-sheet. Due to the high global *B*-factors and presence of diffuse signal, Jackson and colleagues performed three separate *TLS* refinements to model the crystalline disorder (Fig. 3): *Entire molecule*, *Monomer* and *Sub-domain*. All *TLS* refinement attempts improved the *R*_*free*_ values when compared to the standard isotropic *B*-factor refinement; however, there was no significant difference among the final *R*_free_ values from the three *TLS* groupings.

**Figure 3.**
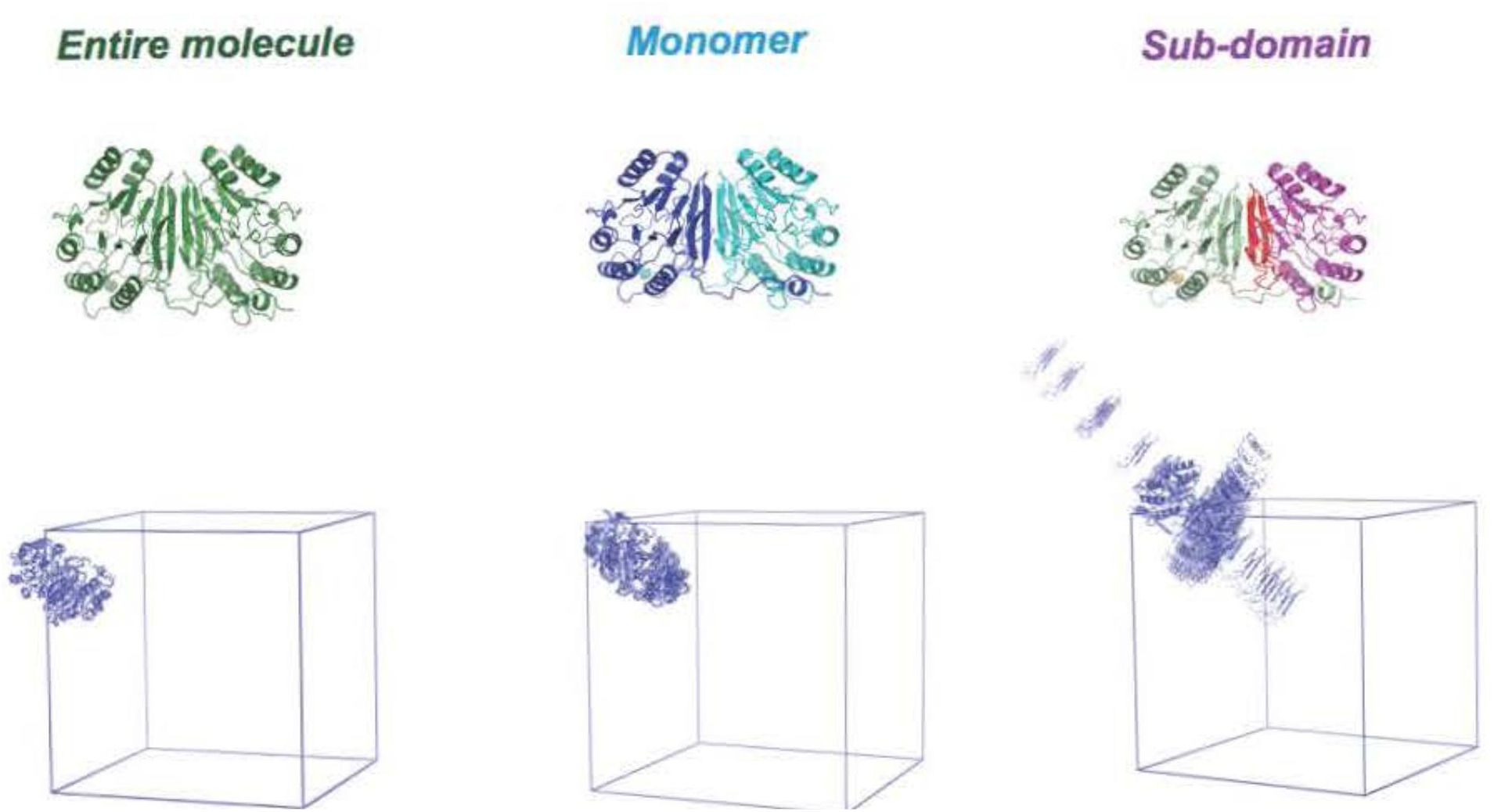
GpdQ TLS ensembles. The GpdQ *TLS* groups are projected onto the protein structure. The corresponding ensembles produced by *phenix.tls_as_xyz* are shown below. Each *TLS* PDB ensemble is shown as a single asymmetric unit outlined by the unit cell. An increase in overall motion is apparent going from left to right. The 20 member ensemble is shown for visual simplicity.

We hypothesized that the diffuse scattering produced by each *TLS* motion would contain significant differences, as diffuse signal is a direct result of correlated motion. The notion that *TLS* refinement produces unique diffuse signal has been suggested previously (Tickle & Moss, 1999). Physical ensembles of the *TLS* motion, rather than an explicit mathematical description, were required to generate 3D diffuse scattering maps from *phenix.diffuse*. Visual inspection confirmed that the ensemble produced by *phenix.tls_as_xyz* replicated the anisotropic motion predicted by *TLS* thermal ellipsoids (Fig. 4). Additionally, we calculated the structure factors predicted by the original *TLS* refinement *Entire molecule* and compared them to *F*_*model*_ values (for example as defined in Afonine et al., 2012) produced by various *phenix.tls_as_xyz* ensemble sizes. The structure factors converged to a global correlation value of 0.965, demonstrating that *phenix.tls_as_xyz* ensembles accurately represent the motions predicted by *TLS* refinement. Physical representation of the underlying motion also revealed that, while two of the *TLS* refinements produced motion with small variances (a necessity within *TLS* theory), the *Subdomain TLS* grouping produced fluctuations that are clearly chemically unreasonable (Fig. 3). Thus, viewing *TLS* refinement in the form of a structural ensemble is a valuable check of the validity of the results, as matrix elements that satisfy the previously described conditions may still produce motions that are clearly implausible.

**Figure 4.**
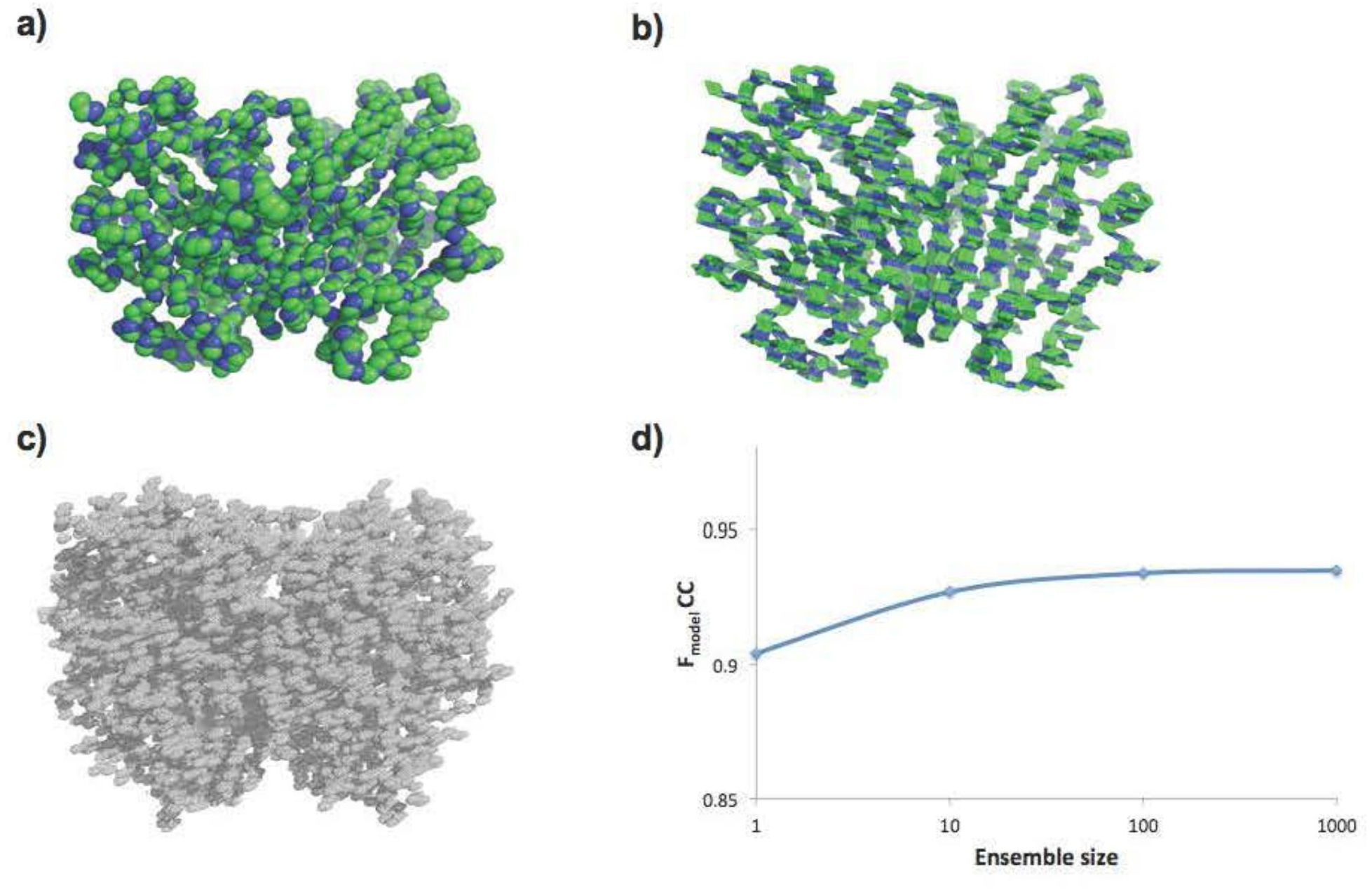
*phenix.tls_as_xyz* ensembles replicate *TLS* anisotropic motion. a) GpdQ backbone with thermal ellipsoid representation of “entire molecule” *TLS* anisotropic *B*-factors. b) *phenix.tls_as_xyz* ensemble backbones produced from “entire molecule” *TLS* refinement. c)Complete electron density predicted by “entire molecule” *TLS* refinement. d) Global correlation coefficient between experimental structure factor amplitudes *F*_*obs*_ of the original GpdQ ‘entire motion’ refinement and *phenix.tls_as_xyz* ensembles of various sizes. Convergence values plateau at 0.935.

## 8. Discussion

While our previous review on the subject (Urzhumtsev et al. 2013) describes the calculation of *TLS* matrices from a known set of vibration and libration parameters, this work focuses on the opposite problem of extracting vibration and libration parameters from a given set of *TLS* matrices. The problem is not as simple as it may at first seem because identical motions may be represented by different vibration-libration combinations. As a consequence, the matrix *S* is not uniquely defined. However, there remains a necessary consistency between the *S* matrix and the *T* and *L* matrix elements. Because current structure refinement programs ignore this constraint, some definitions of *S* may result in *TLS* matrices that cannot be interpreted in terms of physically meaningful motions.

The detailed algorithm for obtaining molecular vibration and libration parameters makes decomposition of the *TLS* matrices into ensemble models possible. The constraints on the matrices can also be used to identify ‘non physical’ combinations of *TLS* matrices. Beyond the well-known conditions of non-negativity for the eigenvalues of *T* and *L*, we also discuss the conditions that relate matrices, which is crucial for ensuring that the *TLS* refinement corresponds to physically meaningful motions. Table 1 sheds more light on how well these conditions are verified for the full collection of PDB structures. For approximiately 4,500 *TLS* sets (roughly 700 PDB entries), correcting the diagonal elements of the *S* matrix as described previously (instead of the standard requirement making its trace equal to zero) corrects the underlying problems. However, even after this correction is applied, however, there still remain a significant number of PDB entries with unphysical TLS matrices.

We suggest that the problem of obtaining physically meaningless *TLS* matrices and the need for their post-refinement correction may be entirely eliminated if *TLS* refinement were to use vibration and libration parameters directly as refined variables. The mathematical basis and computation algorithms for such a refinement is under active development and will be discussed elsewhere. However, eliminating physically meaningless rigid body motions may increase the *R*-factors for some structures. The procedures for analysis and validation of *TLS* parameters, as well as the algorithm for generating a set of models from given libration and vibration parameters, are implemented in the *Phenix* suite as *phenix.tls_analysis* and *phenix.tls_as_xyz*, respectively and are available starting with version dev-1890.

## Acknowledgements

PVA and PDA thank the NIH (grant GM063210) and the *Phenix* Industrial Consortium for support of the *Phenix* project. J. S. F. is a Searle Scholar, a Pew Scholar, and a Packard Fellow. Work in the lab of J. S. F. is supported by NIH OD009180, GM110580, and NSF STC-1231306. This work was supported in part by the Program Breakthrough Biomedical Research, which is partially funded by the Sandler Foundation, and by the US Department of Energy under Contract No. DE-AC02-05CH11231. AU thanks the French Infrastructure for Integrated Structural Biology (FRISBI) ANR-10-INSB-05-01 and Instruct, part of the European Strategy Forum on Research Infrastructures (ESFRI) and supported by national member subscription.

